# Computational synthesis of cortical dendritic morphologies

**DOI:** 10.1101/2020.04.15.040410

**Authors:** Lida Kanari, Hugo Dictus, Athanassia Chalimourda, Werner Van Geit, Benoit Coste, Julian Shillcock, Kathryn Hess, Henry Markram

## Abstract

Neuronal morphologies provide the foundation for the electrical behavior of neurons, the connectomes they form, and the dynamical properties of the brain. Comprehensive neuron models are essential for defining cell types, discerning their functional roles and investigating structural alterations associated with diseased brain states. Recently, we introduced a topological descriptor that reliably categorizes dendritic morphologies. We apply this descriptor to digitally synthesize dendrites to address the challenge of insufficient biological reconstructions. The synthesized cortical dendrites are statistically indistinguishable from the corresponding reconstructed dendrites in terms of morpho-electrical properties and connectivity. This topology-guided synthesis enables the rapid digital reconstruction of entire brain regions from relatively few reference cells, thereby allowing the investigation of links between neuronal morphologies and brain function across different spatio-temporal scales. We synthesized cortical networks based on structural alterations of dendrites associated with medical conditions and revealed principles linking branching properties to the structure of large-scale networks.

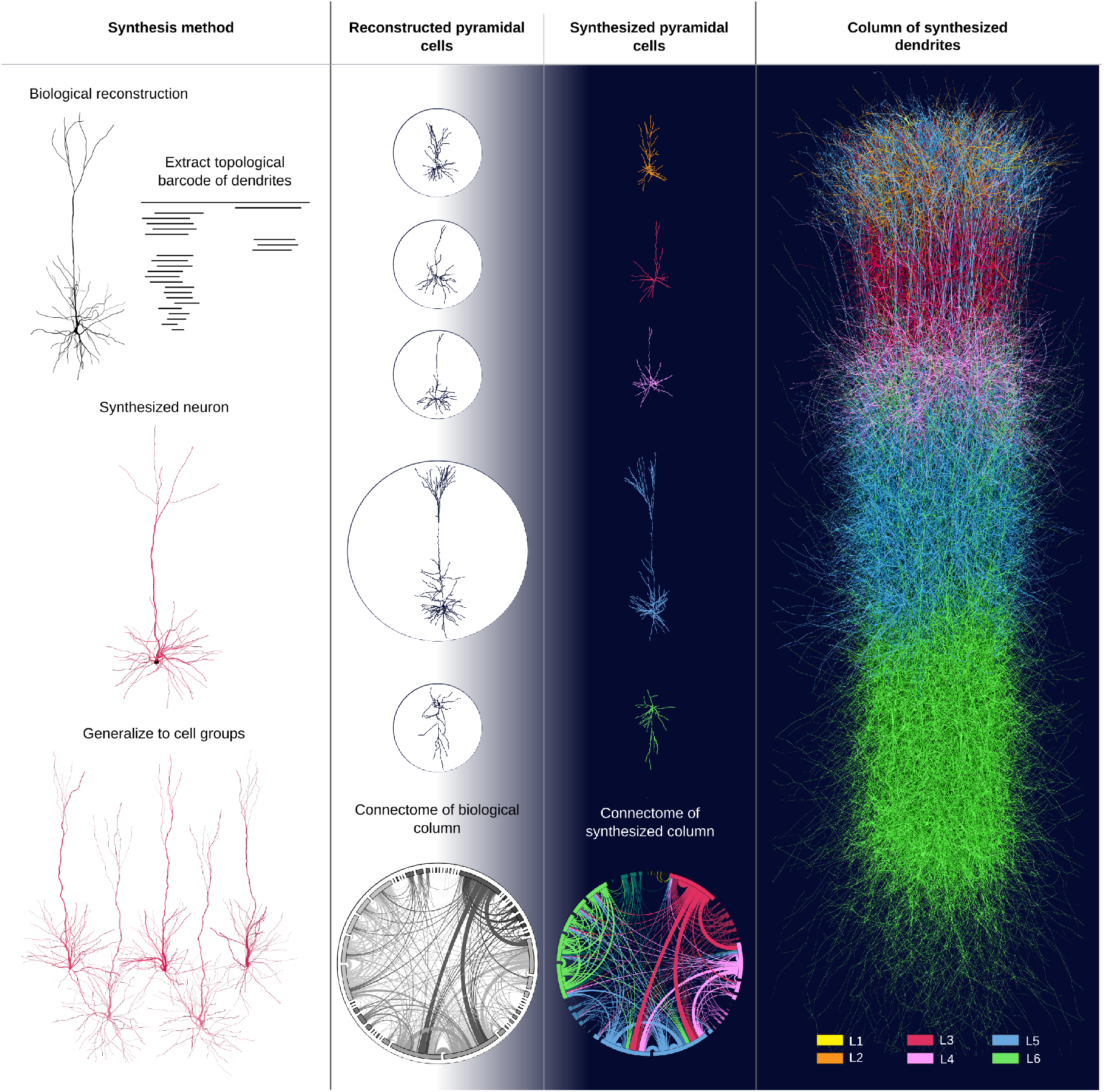
Graphical abstract. A topological model of neuronal shapes is used to investigate the link between the branching patterns of dendritic morphologies and the connectivity of the neuronal networks they form. Starting from reconstructed cells (in black) of cortical dendrites, we extract the topological barcode that is used to create a statistically similar synthesized pyramidal cell (in red), and respectively a group of pyramidal cells of the same morphological type. From reconstructed cells examples of all layers and morphological types we generate synthesized dendrites and build a synthesized cortical column (colors corresponds to cortical layers). The synthesized dedrites are statistically similar to the reconstructed dendrites in terms of morpho-electrical properties and the connectome of the synthesized column (colored connectome) is almost indistinguishable from the connectome of the reconstructed column (greyscale).

## Introduction

Neuronal morphologies play a crucial role in governing a neuronal network’s dynamical properties, as they influence the functions of single neurons (Häusser et al. 2000, Smith et al. 2013, Yi et al. 2017) and constrain the contact points between neurons (Chklovskii 2004, Wen et al. 2009, Cuntz et al. 2012). The importance of structural properties of neurons has been shown in studies that revolutionized our understanding of brain functions, such as the first description of neuronal networks by Ramon y Cajal (Cajal 1899), who argued that the shape of neurons reflects the communication between them, the description of the integrative functions of neuronal dendrites by Willfrid Rall (Rall 1959), and the mathematical modeling of ionic mechanisms underlying the initiation and propagation of action potentials by Alan Hodgkin and Andrew Huxley (Hodgkin and Huxley 1952). Ramon y Cajal argued that the shape of neurons reflects the communication between them (Cajal 1911). A century later, the evidence that the wide variety of neuronal shapes supports the composite functional roles of different cell types is irrefutable, leading to broad a plethora of projects dedicated to harvesting cellular morphologies from various brain regions and species (Peng et al. 2015, Economo et al. 2016, Benavides-Piccione et al. 2019, Gouwens et al. 2019). A burning challenge of our time is to untangle the still largely unknown roles of distinct morphological cell types.

Recent advances in manual (Economo et al. 2016, Benavides-Piccione et al. 2019, Gouwens et al. 2019) and automatic reconstruction techniques (Peng et al. 2015), as well as the systematic collection and registration of digital morphologies in standardized databases (Ascoli et al. 2007, Halavi et al. 2008, Akram et al. 2008), have accelerated the acquisition of neuronal reconstructions. We are still far, however, from having sufficient reconstructions of unique morphologies to populate biologically realistic networks of a whole brain region (1M neurons for the mouse somatosensory cortex, 10M neurons for the mouse isocortex, Eroe et al. 2018, Herculano-Huzel 2006). Digital reconstruction of physiologically realistic neuronal networks (Markram et al. 2015, Egger et al. 2014) requires a large number of distinct neuronal shapes (Shillcock et al. 2016, Landau et al. 2016, Ramaswamy et al. 2012); the reconstruction of a single cell is an expensive and tedious process requiring the collaboration of several reconstruction experts (Farhoodi1 et al. 2019). Even laboratories that dedicate their efforts to large-scale data harvesting can reconstruct only a limited number of cells per year (Janelia, Winnubst et al. 2019, AllenBrain, Gouwens et al. 2019). These datasets are invaluable for identifying the fundamental morphological characteristics of different cell types, but it is unrealistic to expect that adequate numbers of single cell reconstructions can be acquired in this way. A generative model for digital cells that accurately reproduces the shapes of reconstructed neuronal morphologies is thus essential.

A crucial obstacle to the computational generation of neurons (Hillman 1979) is the difficulty in capturing and recreating the correlations between morphological features that arise from highly complex developmental processes, especially given the small number of available examples for each morphological type (m-type). Biophysically accurate models simulate detailed neural growth by integrating the known molecular mechanisms of neuronal development (Zubler and Douglas 2009), thus capturing the correlations in cellular growth. These models focus on the microscopic scale of growth, thereby hindering the computational synthesis of large numbers of neurons. Alternatively, phenomenological synthesis models are based on either fundamental mathematical principles (Luczak 2006, Cuntz et al. 2010) or statistical sampling of the morphological distributions (Ascoli et al. 2001, Koene et al. 2009). Mathematical models study the effect of different factors on neuronal growth, such as spatial embedding (Luczak 2006, Luczak 2010), minimization of wiring cost (Cuntz 2010), and self-referential forces (Samsonovich and Ascoli 2003, Memelli 2013). These models require few parameters and provide good intuition about mechanisms involved in neuronal growth, but require adjustments for different cell types and include additional structural properties. Statistical models (Ascoli et al. 2001, Koene et al. 2009, Lopez-Cruz et al. 2011) generate cells of a given morphological type with high accuracy (Koene et al. 2009), but often overlook biologically relevant feature correlations. In order to identify these feature correlations, manual selection of feature dependencies is required (Lopez-Cruz et al. 2011), which renders statistical approaches computationally expensive and hard to generalize.

The limitations of these models demonstrate the necessity of combining mathematical and statistical properties into a unified synthesis model that circumvents the explicit selection of correlated features, while also being computationally tractable. We developed such a synthesis algorithm based on the Topological Morphology Descriptor, *TMD* (Kanari et al. 2018), which encodes both the topological and the geometric properties of neurons into a single descriptor, the persistence barcode. The persistence barcode of a neuron encodes the start and end distances (i.e., radial or path distance) of all branches within a tree as pairs of numbers. Equivalently, the persistence diagram represents the pairs of start - end distances in a 2*D* plane. We used this descriptor to define the bifurcation and termination probabilities during synthesis; the coupling of these probabilities provides a method to implicitly reproduce key correlations between morphological features. The persistence barcode of different cell types is thus sufficient to capture the different growth mechanisms that lead to the distinct shapes of dendrites. Additional morphological features (soma size, trunk orientation, and thickness of branches, see SI:Synthesis Input) are necessary to capture structures that are complementary to the branching properties. Using the topological neuronal synthesis (TNS) algorithm, we can efficiently synthesize millions of unique neuronal morphologies (10M cells in 4h). A multi-stage validation is then performed to ensure that three modalities of reconstructed neurons are accurately reproduced: morphological characteristics, electrical activity of single cells, and the connectivity of the network they form.

The TNS algorithm is also sufficiently versatile to be applied to a large variety of cell types without parameter optimization. Few exemplar morphologies (for example, for L4 TPC only 15 cells) are required to capture the diversity of neuronal shapes within a reconstructed population. The universality of TNS offers a unique opportunity to model and study systems that are not accessible with current methods. As an example, we generalize the available neuronal reconstructions of the somatosensory cortex (morphological reconstructions published in Markram et al. 2015) to other regions of the neocortex of varying cortical thicknesses. We demonstrate that an appropriate mathematical transformation that accounts for the different neocortical thicknesses is necessary and sufficient to generate pyramidal cells with a gradient of structural properties. This structural gradient of pyramidal cell sizes is in agreement with recent findings, and has been reported to contribute to the unique information processing of neurons depending on their location in the rodent neocortex (Fletcher and Williams 2019).

Last but not least, TNS provides a tool to directly investigate the link between local morphological properties and the connectivity of the neuronal network they form. This approach is of particular interest for medical applications, as it enables the investigation of diseases in terms of the emergence of global network pathology from local structural changes in neuron morphologies. Moreover, TNS offers a technique to generate neuronal morphologies for which very few or no reconstructions are available, as long as an appropriate mathematical transformation can be defined between the target neuronal population and an existing set of neuronal reconstructions. As an example of this process, we reproduced the effects of dendritic alterations associated to stress disorders (Curran et al. 2017, Dioli et al. 2019, Tornese et al. 2019, Sandini et al. 2020). Two relevant structural changes are simulated (Curran et al. 2017, Dioli et al. 2019, Tornese et al. 2019): the shrinkage of dendritic processes and the loss of dendritic branches, by applying two types of mathematical transformations to all the dendrites within a rodent cortical column. Surprisingly, these two types of local dendritic alterations affect the resulting cortical networks quite differently. Neuronal networks formed from dendrites that are gradually shortened collapse rapidly, as they lose connections almost linearly with the total loss of dendritic extent. On the other hand, networks based on dendrites that gradually lose branches of increasing lengths are more resistant to loss of connectivity. These observations suggest the existence of a homeostatic mechanism that preserves the overall network functionality, as long as total dendritic extents remain within a reasonable limit, making the brain networks sufficiently robust to support the complex cortical processes that are fundamental for a healthy brain.

## Results

The morphological development of neurons in the brain is a complicated process that depends on both genetic and environmental components. The processes that contribute to neuronal growth differ between species, brain regions, and morphological types. Advances in experiments and mathematical and computational models have converged on a set of commonly accepted stages of morphological growth: the initiation of neurites, neurite elongation, axon path-finding, and neurite branching (Cuntz et al. 2006, Graham and van Ooyen 2006). These growth stages are useful for computational modeling of the generation of synthesized neurons. In this study we focus on the computational synthesis of dendrites and thus will not consider axon path-finding. While biological development is not simulated, biological principles of morphological growth inform the design of our computational algorithm that synthesizes dendritic morphologies.

### Dendritic synthesis algorithm

The TNS algorithm consists of three main components (Figure 1A): the initiation of dendrites on the soma (see STARmethods, I. Initiation of neurites), branching (see STARmethods, II. Bifurcation / Termination), and elongation (see STARmethods III. Elongation of neurite) of neurites. First, the neuronal soma is generated based on a radius sampled from a biological distribution. Then, the number of neurites is sampled from the biological distribution of the corresponding cell type. Each neurite is assigned an initial orientation and a barcode, based on the reconstructed neurites of the respective morphological cell type.

**Figure 1:**
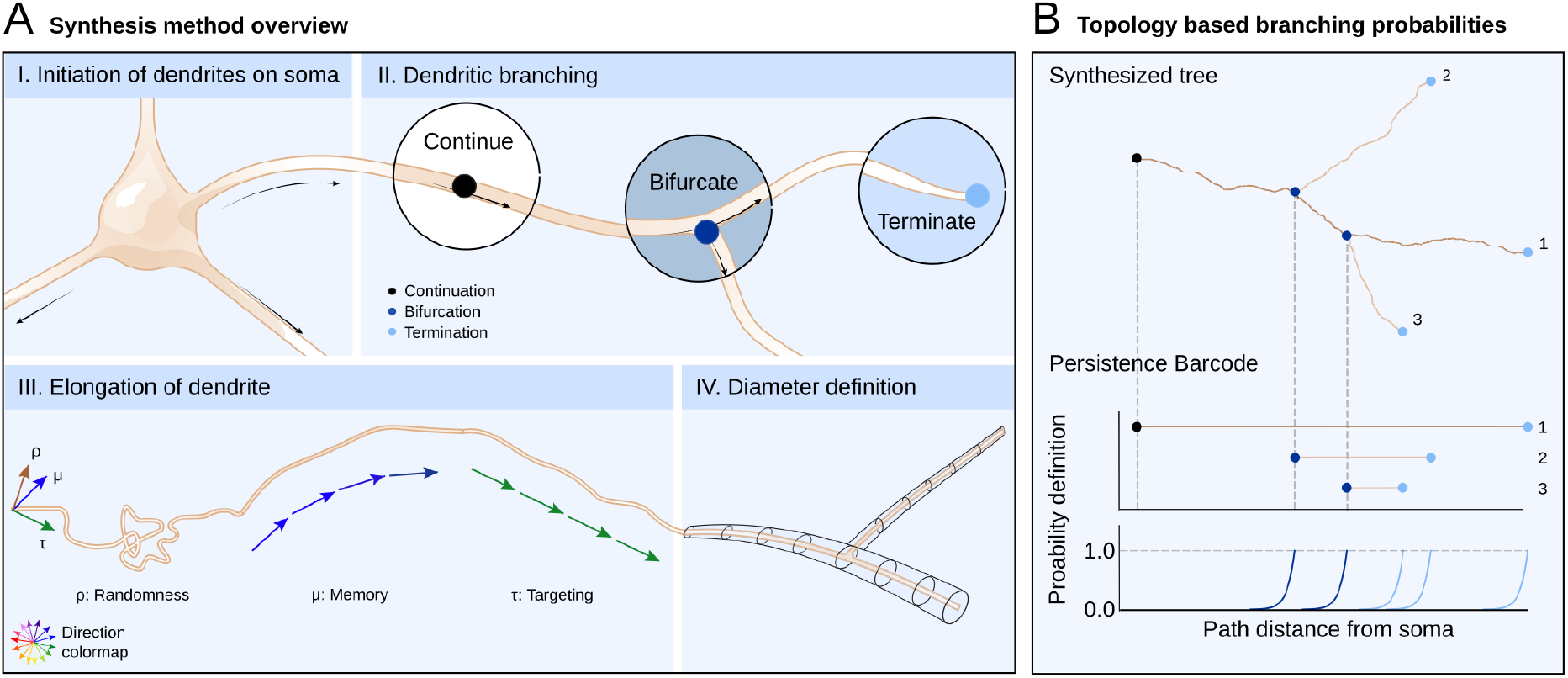
Method of Topological Neuron Synthesis. A. Overview of dendritic synthesis based on four stages of growth. I. Soma generation and initiation of the dendrites on the soma surface. II. Stochastic definition of bifurcation, termination, and continuation based on topological descriptor (B). III. Dendritic elongation: during continuation the branch grows based on a segment length and direction. The direction is chosen as a combination of three parameters: randomness, memory (based on the previous directions within a branch), and targeting (based on the initial direction of a branch). IV. Diameter definition, as a final step, is based on the biological distributions and is subsequent to the branching steps. B. Branching based on the topological morphology descriptor (TMD) of a neuronal tree: the probability to bifurcate, terminate, or continue depends on the path distance from the soma and the joint probabilities derived from the TMD of a neuronal morphology. The start of a bar in the TMD increases the bifurcation probability; the end of the bar increases the termination probability. Note that each new bar has to be smaller that its parent, so the growth is performed from larger to smaller branches.

Subsequent steps of the growth take place in a loop. Each branch of the tree is elongated step by step, as a combination of the following components: a random direction, *ρ*, the target orientation, *τ*, and a memory of the previous steps, *μ* (Figure 1A-III, see STARmethods III. Elongation of neurite). At each step, the growing tip is assigned probabilities to bifurcate, to terminate, or to continue that depend on the path distance from the soma and are defined by the bars of the sampled barcode (Figure 1A-II, see STARmethods, II. Bifurcation / Termination). Each bar within the barcode can only be used once. The growth terminates when all the bars of the input barcode have been used. Finally, as an independent step, the diameters of the tree are assigned based on diameter distributions sampled from the reconstructed cells (Figure 1A-IV, see STARmethods IV. Generation of tree tapering). The details of the algorithm are described at the STARmethods section.

The three main modalities that we want to reproduce with synthesized cells are morphology (Scorcioni et al. 2008), electrophysiology (Van Geit et al. 2016), and connectivity (Van Pelt et al. 2010, Van Pelt and Van Ooyen 2013). In order to ensure that the synthesized cells recreate all three properties, the microcircuit that was published by Markram et al, 2015 was used as reference. This microcircuit was built based on reconstructions of juvenile rat neurons across all cortical layers for a variety of morphological types. First, we ensure that the morphological and electrical properties of the reconstructed cells are reproduced by neurons whose dendrites have been synthesized with the TNS algorithm. Then, a set of synthesized cells of an m-type is compared against the set of reconstructed cells of the same m-type. Finally, a network of cells is generated based on the neurons with synthesized dendrites, and compared to the connectivity of the original network.

## Single neuron synthesis

There are two major types of cortical neurons, based on their functional roles: excitatory and inhibitory. Excitation is mainly mediated by pyramidal cells, with the exception of the spiny stellate cells (L4), and use glutamate as a neurotransmitter. Inhibition is mediated by interneurons, which use GABA as a neurotransmitter. The TNS algorithm is used for the computational synthesis of dendrites of both interneurons and pyramidal cells of a large variety of morphological types. Because these two major cell classes present distinct morphological properties, synthesized pyramidal cells and interneurons are presented independently (Figure 2A-B). Interneurons have only basal dendrites, i.e., dendrites that emanate from the base of the cell body and are localized mainly around the soma, are less complex than pyramidal cells, which also have apical dendrites that reach to higher cortical layers, typically ascending towards the pia, and present a wider diversity of shapes.

**Figure 2:**
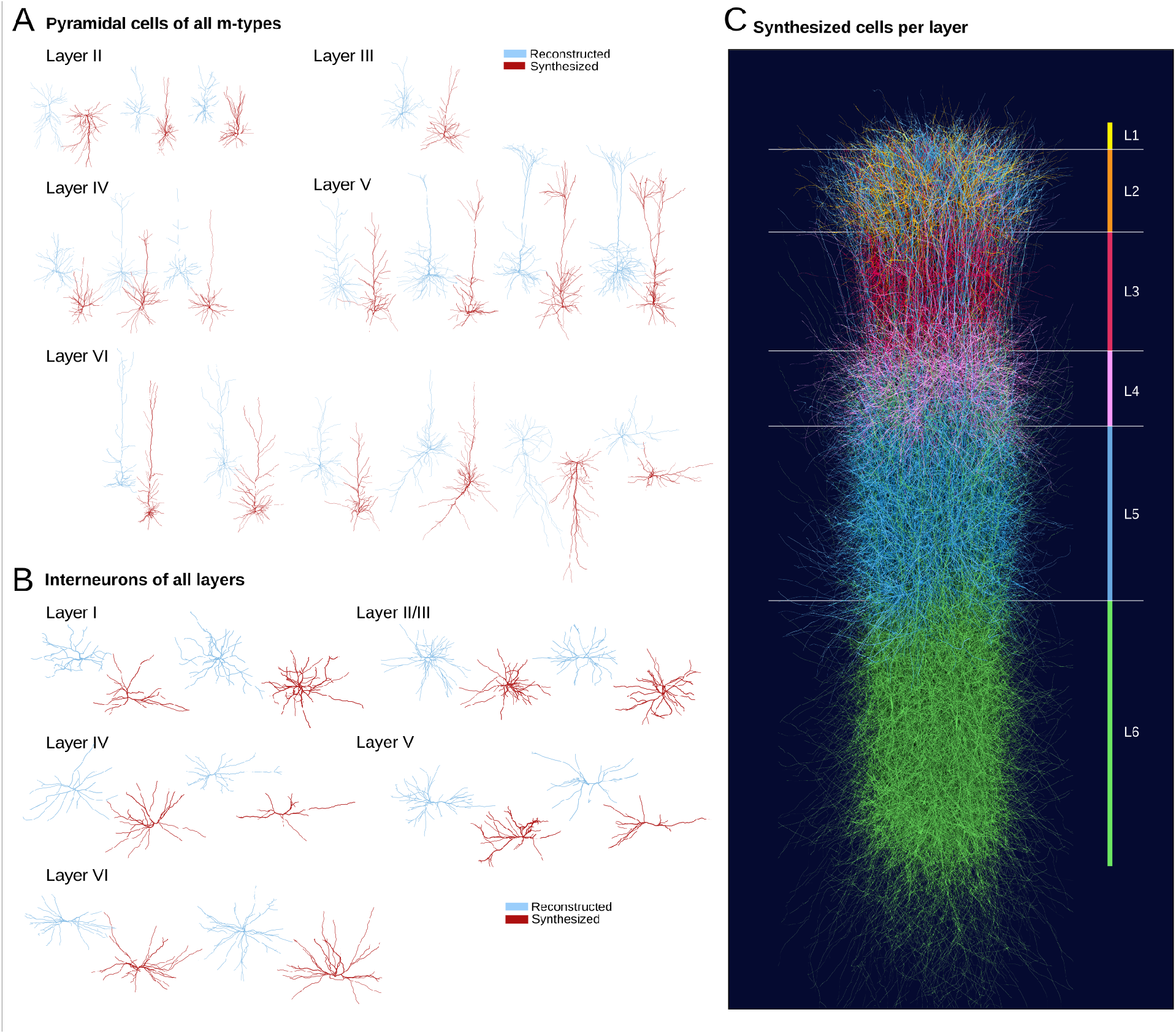
Comparison of reconstructed and synthesized dendritic shapes. A. Reconstructed (blue) and synthesized (red) pyramidal cell dendrites of all rodent cortical m-types from layers 2 to 6. B. Reconstructed (blue) and synthesized (red) dendrites of rodent cortical interneurons of layers 1 to 6. Not all interneuron morphology types are reported, as they differ mainly in their axonal branches and not significantly on the basal dendrites, as illustrated. C. A cortical column of synthesized dendrites of all layers, colors correspond to cortical layers from 1 to 6.

### Reproducing characteristic neuronal shapes

Interneurons represent about 15% (Gonchar et al. 2006, Lefort et al. 2009) of neuronal cells in the cortex. There are many interneuron types, which are distinguished mainly by their axonal shapes, as they do not vary greatly in their basal dendrites. A subset of reconstructed interneuron dendrites of different layers is presented in Figure 2C, (top, blue). Computationally synthesized dendrites of the same layers (Figure 2C, (bottom, red) are generated based on the corresponding topological profiles of the biological reconstructions. The TNS algorithm generates basal dendrites that reproduce the characteristic shapes of different morphological interneuron types.

Pyramidal cells (PCs) represent the majority of neurons in the cortex (≈ 85%, Gonchar et al. 2006, Lefort et al. 2009). The wide variety of apical dendrite shapes imparts unique functional properties to PCs and forms the basis for integrating signal inputs from different cortical layers (Larkum et al. 2007, 2009, Spruston 2008). In a previous publication (Kanari et al. 2019), we distinguished 18 pyramidal cell types, 17 of which could be objectively supported. In Figure 2A we present examples of reconstructed morphologies (in blue) of the 18 different m-types of pyramidal cells from layers 2 to 5. Synthesized dendrites based on the topological profiles of the same types of pyramidal cells are illustrated in Figure 2A (in red). Thus, the TNS algorithm generates cells that reproduce the characteristic shapes of the dendrites of distinct morphological pyramidal cell types.

### Reproducing morpho-electrical properties

To ensure that each synthesized cell is consistent with the reconstructed cells, both its morphological and electrical properties need to be validated. A set of 100 synthesized cells was generated, based on the morphological properties (persistence diagram and morphometrics) of a selected reconstruction of a L3 TPC (Figure 3A). First, the topology of the synthesized cells was validated, by comparing the radial-persistence diagram of the reconstructed to the synthesized cells’ (Figure 3B). Subsequently, the morphometrics of reconstructed and synthesized cells were compared (Figure 3C). Finally, the electrical model optimized on a population of L2/3 pyramidal cells (Van Geit et al. 2016) was applied to the reference reconstructed cell and a synthesized cell. Similar to the morphological validation, the Fnorm was used to quantify how well the resulting morpho-electrical combination matches with the statistics of the original experimental data for the 120% threshold current step amplitude (Figure 3F,G,H). Their electrical traces in response to the tested stimuli are presented in Figure 3F-G and the electrical features in Figure 3H. For both electrical and morphological comparisons, each feature (Figure 3C and H) is normalized according to equation 1.

**Figure 3:**
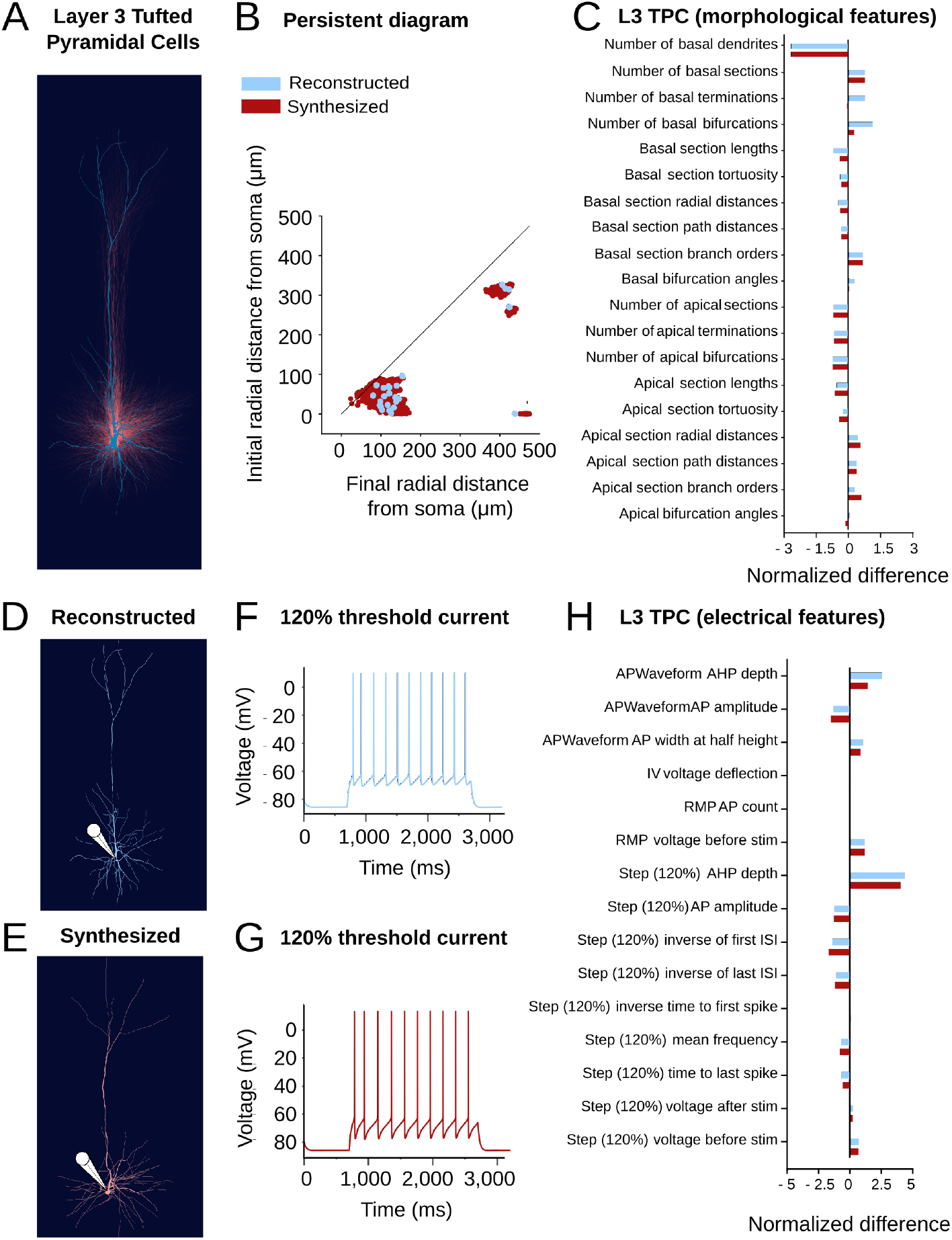
Validation of single cell morpho-electrical properties. A reconstructed layer 3 tufted pyramidal cell (A, blue) is used as input for 100 synthesized L3 TPCs (A, red). B. Comparison of topological persistence diagrams of the reconstructed cell and 100 synthesized cells. C. Comparison of 19 dendritic morphometrics (normalized based on the mean morphological feature values for the L3 TPC population) for a reconstructed and a synthesized cell. The reconstructed (D) and synthesized cell (E) are electrically simulated according to a model optimized on the electrical properties of L3 TPC cells. The electrical response (120% threshold current step) of the reconstructed cell (F) is compared to the synthesized cell’s (G). H. Comparison of 15 electrical properties of dendrites (normalized based on the mean electrical feature values for the L3 TPC population.

Due to the stochastic component of the growth process, the synthesized cells are not identical to the reconstructed cell (Figure 3A&B), thus increasing the morphological diversity. The morphological and electrical properties of the synthesized cells are statistically similar to the respective properties of the reconstruction, as the normalized error is similar for the synthesized and the reconstructed cell for all properties (Figure 3C&H). Overall, the TMD-synthesis algorithm with the selected stochastic parameters (randomness *ρ* ≈ 20%) reproduces the morphological and electrical properties of the reference reconstruction.

## Synthesis of a neuronal population

In order to validate the synthesis algorithm, it is essential to generalize the results of the previous section by using as input a large number of biological reconstructions. Note that the algorithm randomly samples a persistence barcode extracted from the reconstructed population until all dendrites of a cell are grown. We generated synthetic cells for all the rodent cortical cell types reported in previous publications (Markram et al. 2015, Kanari et al. 2019, Marx et al. 2013), using as input a set of reconstructions from the BBP dataset. One hundred cells were computationally synthesized for each pyramidal and interneuron type.

### Reproducing morphological properties

Detailed validations were performed in order to ensure the good quality of the synthesized cells. The synthesized cells of type L5 TPC:A were validated against the reconstructed cells using a large set of morphometrics (Ascoli et al. 2008, Scorcioni et al. 2008), a subset of them is presented in Figure 4 for the apical (top) and basal dendrites (bottom). The statistical distributions of the morphological features (such as number of sections, section lengths, bifurcation angles and branch orders) of the synthesized cells closely match the distributions of the reconstructed cells for both the apical and the basal dendrites.

**Figure 4:**
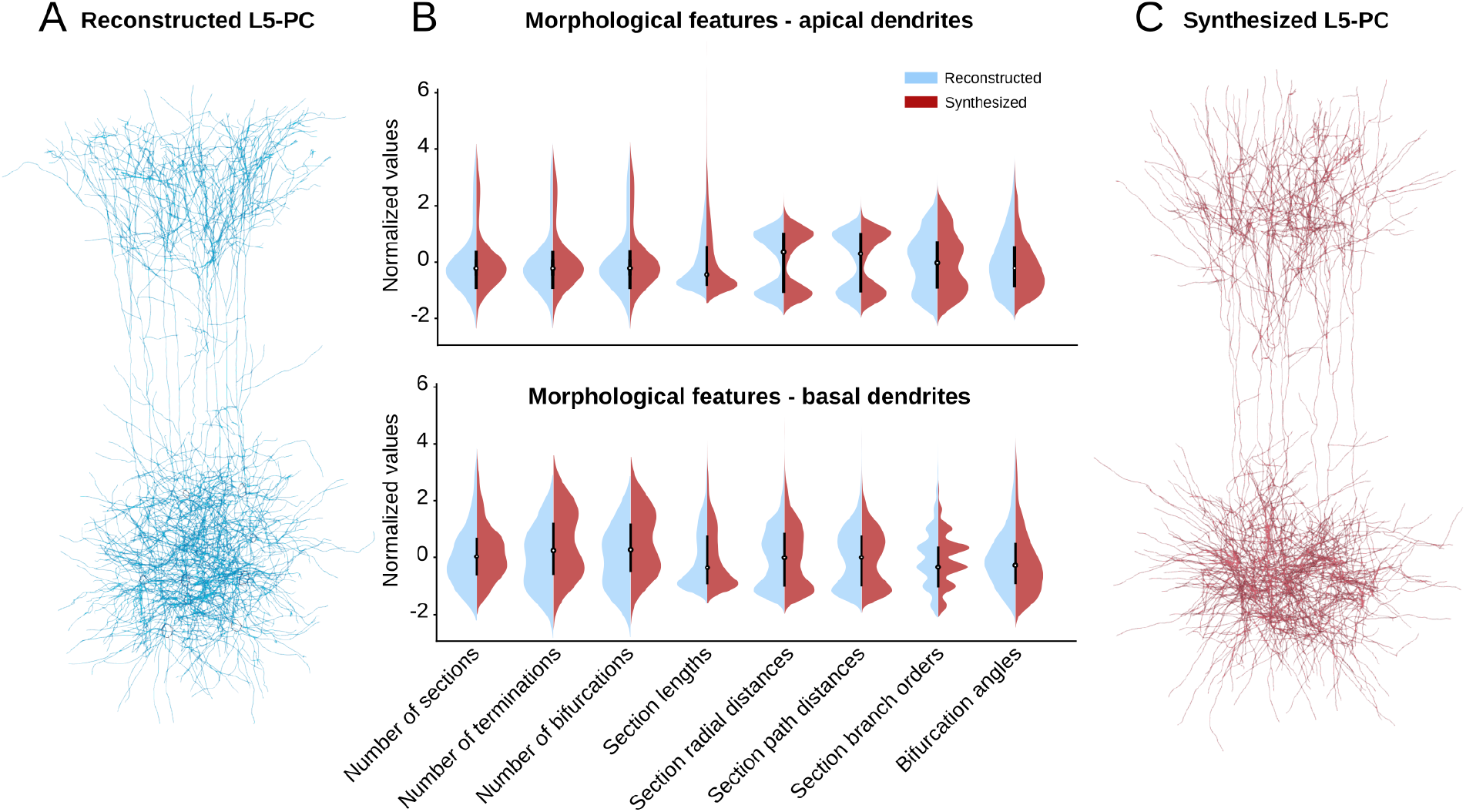
Morphological validation of L5 synthesized pyramidal cells. A set of L5 TPC reconstructions (A, blue 35 cells) is used as input to generate a population of synthesized cells of the same type (C, red, 100 cells). The violin plots of morphological properties for apical (B, top) and basal (B, bottom) dendrites of the reconstructed cell (in blue, left side of violins) and the synthesized cells (in red, right side of violins) are reported. The morphological features of apical dendrites (such as number of sections, section lengths, bifurcation angles and branch orders) are indistinguishable between reconstruction and synthesis populations. Similarly, features of basal dendrites of synthesized cells match the properties of reconstructed basal dendrites. Differences in termination / bifurcation numbers in basals are due to preprocessing of input data to retain only intact parts of the dendrites from incomplete reconstructions.

Artefacts of the reconstruction techniques, as described in C. Synthesis input, that we chose not to reproduce result in minor divergence between specific properties of the two populations. This is due to pre-processing of the data that are used as input to the synthesis algorithm. Namely, the original reconstructed cells have branched of high tortuosity. This is reportedly an effect of the reconstruction process (Conde-Sousa et al. 2016, Farhoodi et al. 2019) that we choose not to reproduce. In addition, only intact branches (i.e., branches that are not cut due to experimental techniques) are used as input in synthesis. As a result, the total number of bifurcations and terminations per basal dendrite is higher for synthesized cells compared to the reconstructed cells’. For apical dendrites the same morphometrics are well reproduced, since no apical trees were excluded from synthesis input. For cell types with five or more available reconstructions, the algorithm generated cells that are statistically close to the input population. For cell types with few available reconstructions (five or fewer), the small number of input cells did not suffice to capture the diversity of the reconstructed cells. We further investigate this aspect in section Versatility of the synthesis algorithm.

### Reproducing morphological correlations

The TNS algorithm generates dendrites that closely approximate the morphometrics of the reconstructed cells, as well as their interdependencies. Correlations between morphological features are reportedly essential for any synthesis method (Lopez-Cruz et al. 2011). However, does TMD-based synthesis implicitly account for correlations, or do we need to define them explicitly? In previous studies, the explicit description of correlated morphometrics was either obtained manually (Koene et al. 2009, Van Pelt and Van Ooyen 2013) or optimized with complex algorithms (Lopez-Cruz et al. 2011). The manual identification of feature correlations is problematic, as experts disagree on the optimal set of features that describes neuronal morphologies (DeFelipe et al. 2013). The set of optimal morphometrics may also not generalize across different cell types. On the other hand, complicated machine learning techniques that infer feature correlations risk over-fitting when only a few reconstructions are available. In this case, instead of capturing the biological principles of neuronal morphologies, the algorithm overestimates local properties, reproducing the noisy properties of input cells.

To investigate whether TMD is important to capture the correlations between morphological features, we synthesized cells that do not take as input the joint probability distribution encoded in the TMD. Instead, the marginal bifurcation and termination probabilities were used as independent parameters (see SI, Figure S6). If bifurcation and termination probabilities in reconstructed cells were in fact independent, this algorithm would suffice to reproduce the branching patterns of the neuronal morphologies (Luczak 2006, Cuntz 2010). Interestingly, in this case the synthesized cells generated by this algorithm were significantly different from the original reconstructions, indicating that the correlations encoded in the persistence barcodes of dendrites are essential to the synthesis algorithm.

### Reproducing morphological diversity

Another challenge in synthesis is the sparsity of input data for many cell types, which makes it difficult to reproduce the morphological diversity of neurons. If few biological reconstructions are available (fewer than five cells), it is not possible to recreate this cell type in its in-vivo conditions. Starting from groups of cells with a large number of available reconstructions, such as pyramidal cells of layers 3-5, we investigated how many cells are required as synthesis input to approximate the morphological diversity of the whole population. Figure 5 illustrates how the number of cells used to define the input data for synthesis affects input (path-distance) and “emergent” (branch order and radial distance) morphological features.

**Figure 5:**
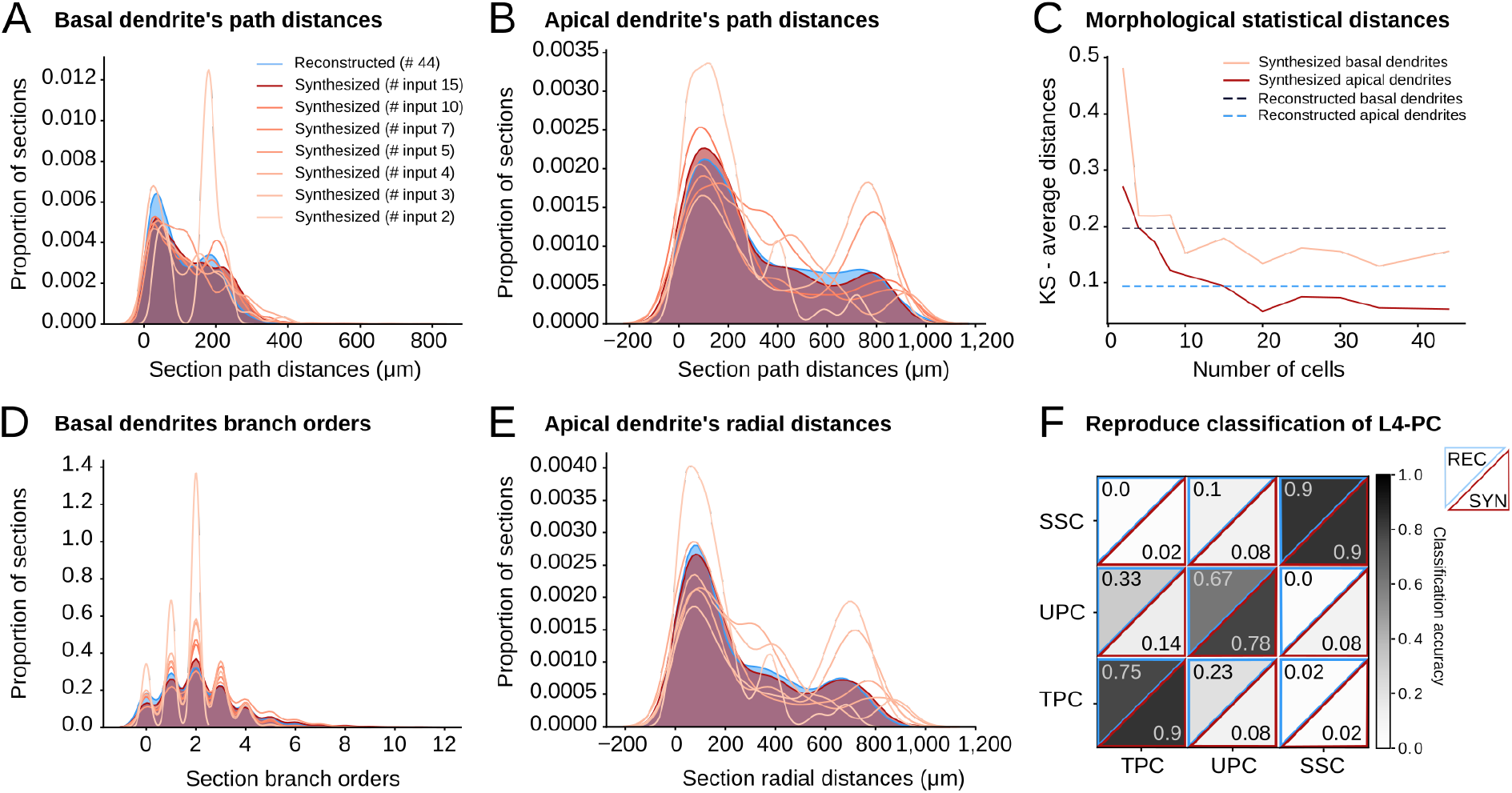
TNS reproduces morphological diversity. Comparison of dendrites from reconstructed L4 TPC cells (in blue, 44 original reconstructions) and synthesized cells (red shades from lighter to darker according to number of randomly sampled neurons used as synthesis input: from 2 to 15). Comparison of path distance (A, input to algorithm) and branch order (B, emergent property) distributions for basal dendrites. Comparison of path distance (D, input to algorithm) and radial distance (E, emergent property) distributions for apical dendrites. The original distributions are approximated with a subset of input cells (15 out of 44). C. Average statistical (Kolmogorov-Smirnov) distance of a set of morphometrics, within reconstructed cells (in blue) and between reconstructed and synthesized cells (in red) with respect to number of randomly sampled neurons used as synthesis input. F. TMD based classification of three L4 PC types for reconstructed (top left, blue) and synthesized (bottom right, red) cells. Classification accuracy is same or higher for the synthesized population.

Varying the number of randomly selected input cells *N* from a population of 44 L4 TPC reconstructed cells, we identified the minimum *N* such that the synthesized population approximates well the morphological diversity of the reconstructed population. While a sample size of *N* ≤ 10 was not sufficient to approximate the diversity of the reconstructed cells with respect to path, radial distances and branch orders, for *N* ≥ 15 (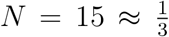 of reconstructions) both input and emerging morphometrics were reproduced well (Figure 5A-D). In addition, for *N* ≥ 15 the overall cumulative error (KS-distance, Figure 5E) of all morphometrics is minimized for both basal and apical features.

Finally, we computed the classification accuracy of the three layer 4 pyramidal cell types (L4 TPC, L4 UPC, L4 SSC)for reconstructed and synthesized cells. The three cell types were synthesized using the same input parameters, and the persistence barcodes of the three respective groups of pyramidal cells. The same classifier (Decision Tree from the sk-learn Python package) was then used to classify the TMDs of both the reconstructed and the synthesized cells into the three original classes, and a leave-one-out accuracy measurement was calculated (Figure 5F). The synthesized cells were classified (Figure 5F, bottom right) into their original classes with at least the same accuracy as the reconstructed cells (Figure 5F, top left).

### Versatility of the synthesis algorithm

As shown above, TMD-based synthesis reproduces the morphological properties of reconstructed cells, while preserving the diversity of cells of a morphological class. However, it is not sufficient to generate cells that reproduce an input population. A simple example to illustrate the limitations of this approach is the growth of neurons within brain regions with varying anatomical properties, such as cortical thickness, which is not constant within the neocortex. Individual animals of the same species and age have variable cortical thickness (De Felipe et al. 2011, Fletcher and Williams 2019). A synthesis algorithm that merely reproduces the original population has serious limitations, as reconstructions are collected from a large number of individuals with varying anatomical brain properties. Statistical and biophysical models are often limited to reproducing the input populations, due to the detailed input that is consumed by the algorithm. On the other hand, mathematical models are adaptable to a variety of different set-ups.

The TNS algorithm can be modified by applying a relevant mathematical transformation to the persistence barcodes. These mathematical transformations include scaling, which alters the size of a dendrite; rotation, which adapts the orientation of the cell; and altering the number of branches of a selected length. To illustrate the versatility of the TNS algorithm, cells for varying cortical thicknesses were synthesized. The persistence barcodes were scaled according to different percentages from 100% to 10% (Figure 6B). We demonstrate that scaling the input barcodes is correctly translated to other morphometrics, as seen in Figure 6 further supporting our argument that feature correlations are accurately represented in the persistence barcodes. For example, the total length of synthesized cells based on scaled barcodes (Figure 6C, D, E) is equivalent to the total length of scaled reconstructions for all layer 4 PCs.

**Figure 6:**
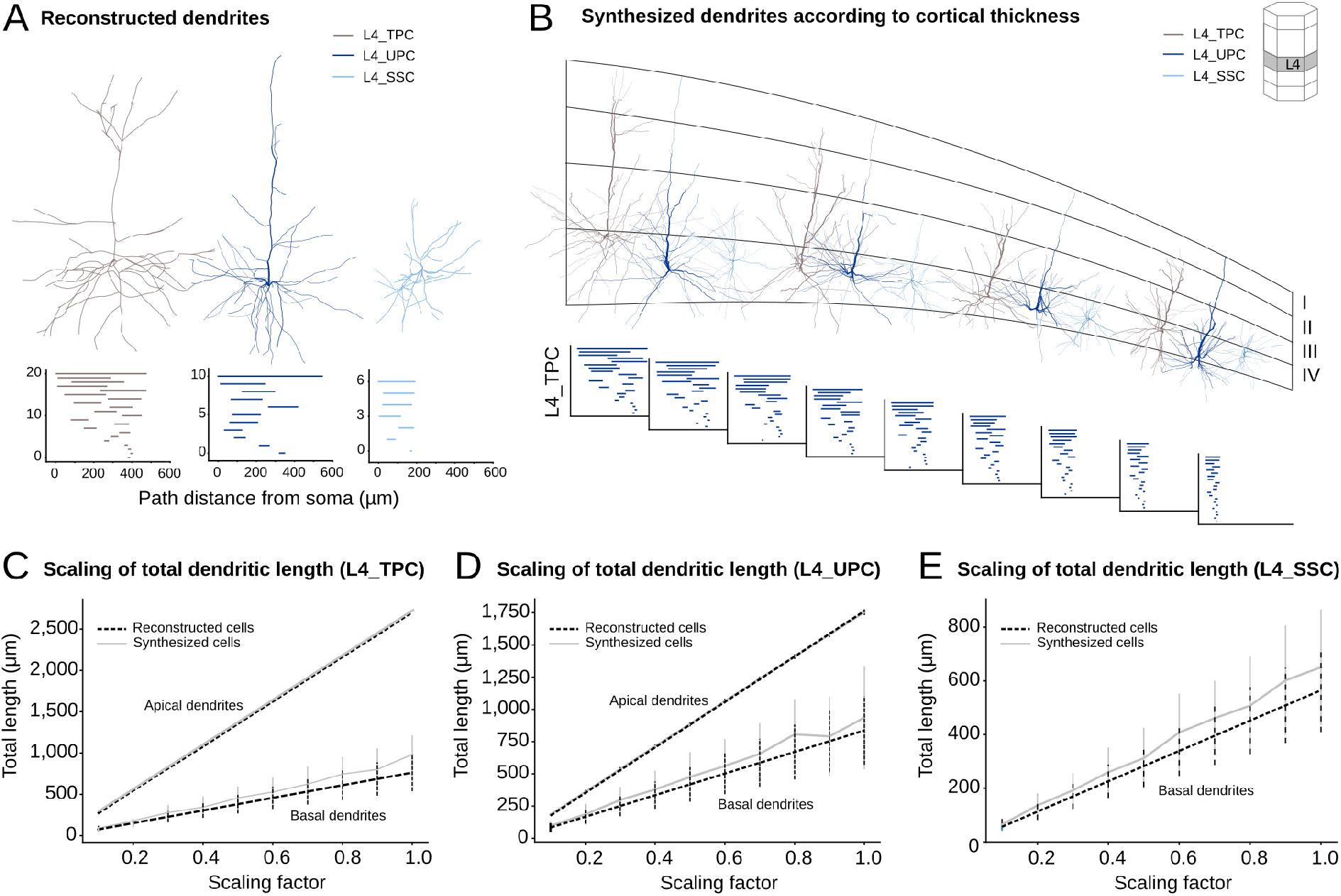
Generalization of topological synthesis for varying cortical thickness. A. Exemplar biological reconstructions of three layer 4 pyramidal cell types: L4 TPC (grey), L4 UPC (deep blue), L4 SSC (light blue) and the corresponding persistence barcodes, used as synthesis input. B. Scaling of input persistence barcodes and resulting synthesized dendrites ((1.0, 0.8, 0.6, 0.5) of original barcodes). The scaled (from 1.0 to 0.2) barcodes of synthesized L4 TPC apicals presented at the bottom. Total dendritic length of layer 4 cells, as a function of shrinkage factor for basal (bottom) and apical (top) dendrites compared to expected values of scaled biological lengths (black dashed, computed as scaling factor multiplied by total length od reconstructed dendrites) and synthesized (grey continuous) dendrites of L4 TPC (C), L4 UPC (D) and L4 SSC (E). Note that L4 SSC do not have apical dendrites even though they are excitatory cells, therefore only basal dendrite statistics are shown.

## Network of synthesized dendrites

### Reproducing the connectivity of a reconstructed microcircuit

In the previous sections, we demonstrated that the topological synthesis generates dendrites that match the morpho-electrical properties of biological reconstructions. In order to verify that synthesized dendrites are suitable for digital simulations of neuronal networks the connections that they form need to be validated. For this reason, we created a replica of the 2015 Markram et al. digital reconstruction of the rat cortical microcircuit, from now on referred to as “reconstructed” microcircuit, built from synthesized dendrites instead of reconstructed cells. Starting from the initial position of each cell, which corresponds to the original position in the reconstructed microcircuit, the dendrites of all neurons were computationally synthesized according to their morphological type. The original axons of the reconstructed microcircuit were used for the definition of the appositions (touch points between dendrites and axons). The connectivity of the synthesized network was then computed (methods described in Markram et al., 2015) and compared to the connectivity of the reconstructed microcircuit.

In Figure 7 we present the statistical properties of the synthesized circuit (B) in comparison to the reconstructed microcircuit (A). The connectome of the microcircuit grouped by m-type (1), the connection probabilities (2), and the numbers of synapses per connection (3) of the synthesized network are in statistical agreement with the reconstructed microcircuit of Markram et al. 2015. Since the initial positions of the cells and the axonal reconstructions of the reconstructed circuit are preserved, we ensure that the synthesis of the dendrites does not significantly alter the statistical properties of the dendrites’ connectivity. The differences of the statistical properties of the two circuits (connectome, connection probability and number of synapses, Figure 7C) are close to zero and significantly lower that the standard deviation of the respective properties.

**Figure 7:**
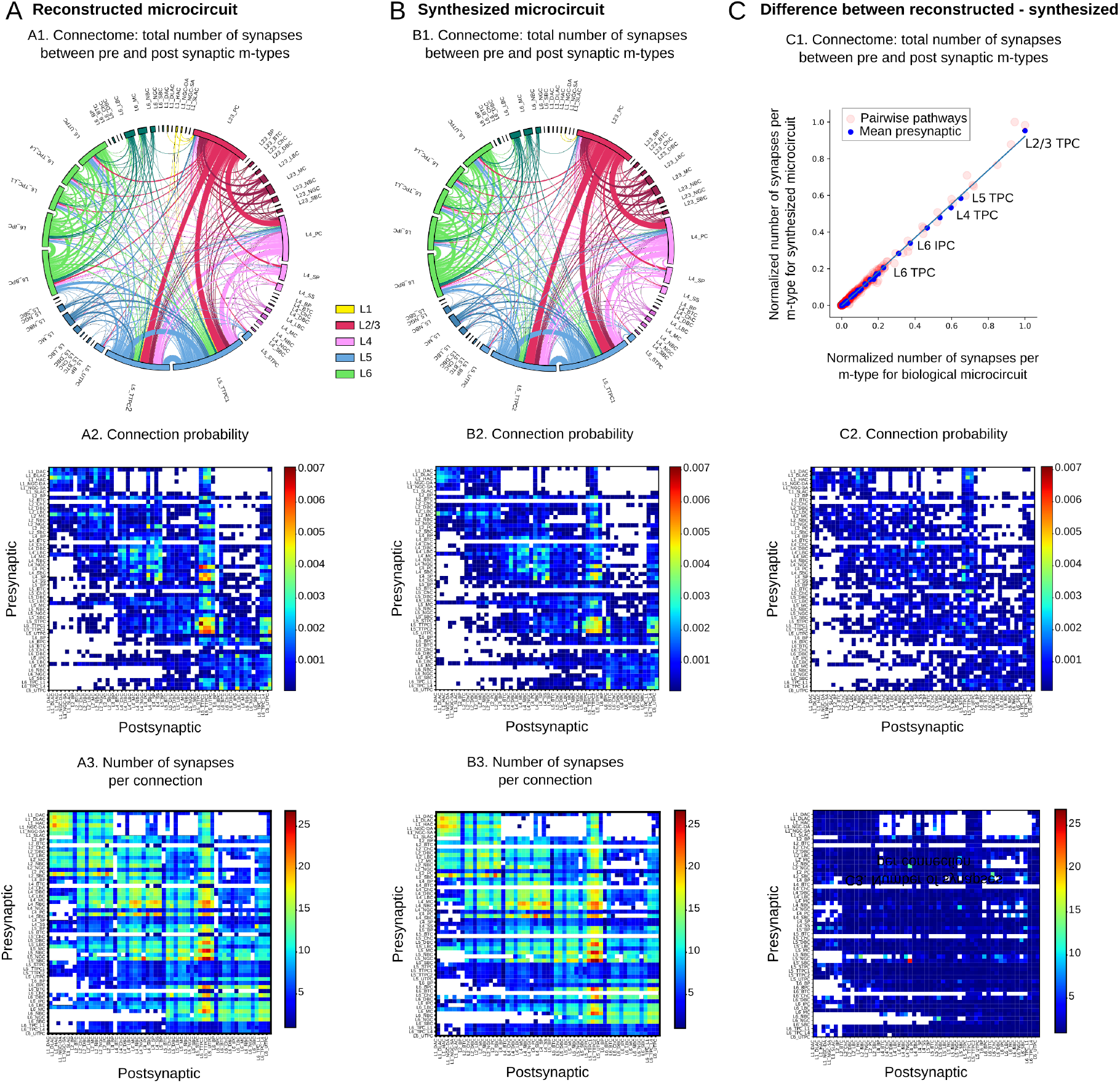
Comparison of connectivity for synthesized and reconstructed networks. A. The connectivity properties of a reconstructed microcircuit (Markram et al. 2015). B. The connectivity properties of a microcircuit of fully synthesized dendrites, and reconstructed axons. C. Difference between reconstructed and synthesized microcircuits. (I) The connectome of the reconstructed microcircuit grouped by m-type. Colors group m-types by layer and correspond to axonal outputs. The thickness of ribbons is proportional to the total number of synapses. (II) Connection probability. A matrix of average connection probability per pathway (350 *μm*, central micro-column, 10K pairs). (III) Synapses per connection. A matrix of the average number of synapses per connection for multi-synapse connections formed between the 55 m-types (10K pairs).

**Figure 8:**
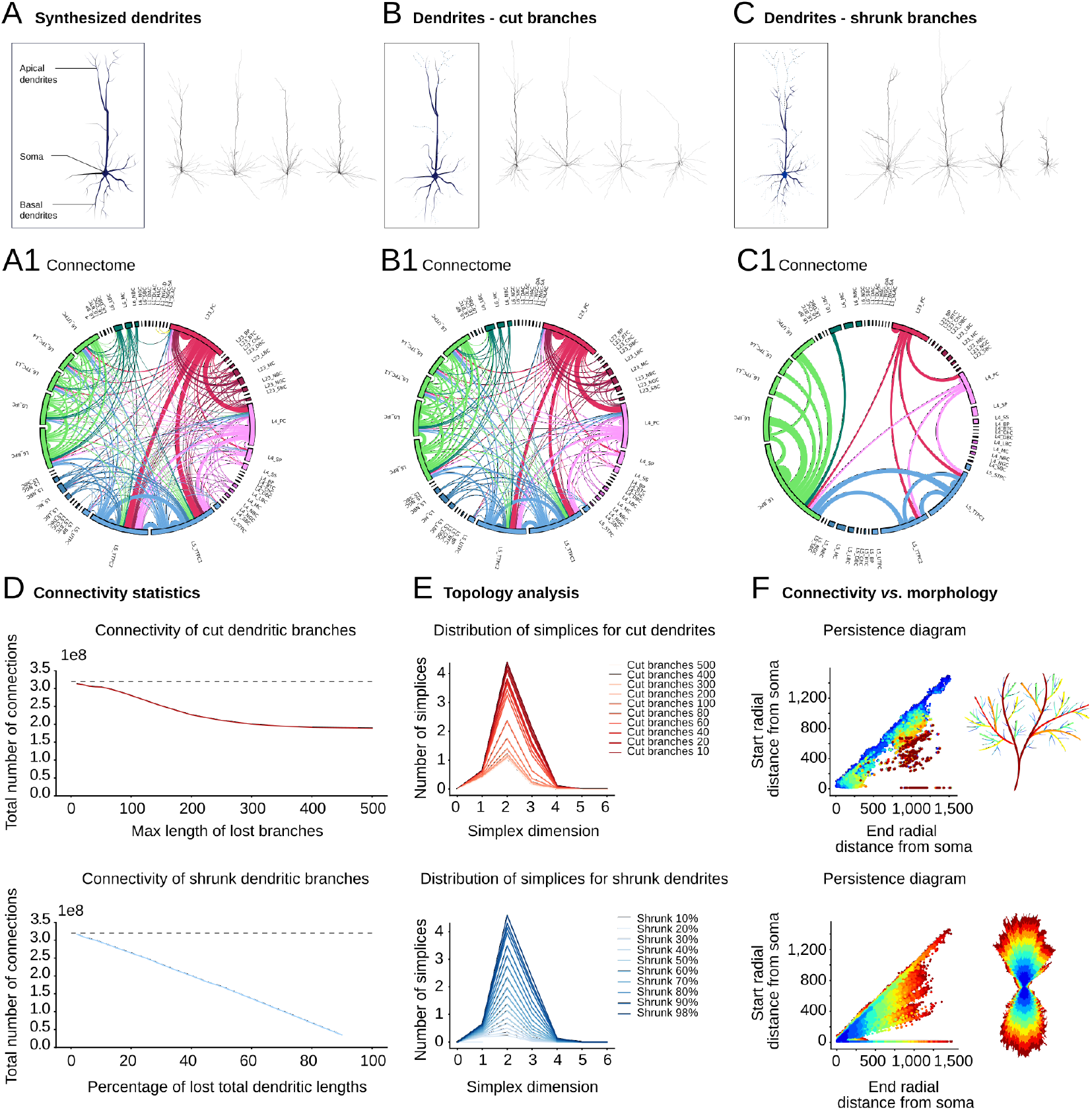
Connectivity of synthesized networks based on structural alterations of dendritic morphologies. Schematic representation and examples of layer 5 synthesized pyramidal cells (A), in comparison to cut dendritic branches (B, lengths above 10, 100, 200, 400*μm*), and shrunk dendrites (C, 98%, 90%, 60%, 30%). Connectome (presented in subpanel 1) of each synthesized microcircuit (A: synthesized, B: cut branches of lengths above 400*μm*, C: shrunk dendrites 10%) (connections above 150K shown, line thickness corresponds to connection strength). D. Total number of connections for alterations of type B (red) and C (blue) compared to synthesized network A (black).E. Topological analysis of corresponding networks; distribution of directed simplices for alterations of type B (red, top) and C (blue, bottom). F. Morphological characteristics and connectivity, with respect to alterations of type B (top) and C (bottom). The main branches form the majority of connections (top) and larger dendritic extents (bottom) form more connections. Colormap corresponds to normalized number of connections: from maximum number of connections (3.5 × 10^8^ in red) to minimum (10^7^ in blue).

### Medical applications

As novel techniques (Watts et al. 2013, Sarifi 2013) emerge for the treatment of and analysis of brain disorders (Meng et al. 2018, van den Heuvel and Sporns 2019), the need to find new ways to predict their outcome becomes increasingly imperative. These techniques are frequently applied to animal subjects (Meng et al. 2018). It is therefore important to find a way to predict how the effects of these drugs, which are tested on smaller mammalian subjects, generalize to primates. The accurate prediction of the effects of complex treatments on humans requires accurate and detailed models of neuronal networks. We propose the first step in this direction. The TNS algorithm enables us to generate synthetic neuronal morphologies for which very few or no reconstructions are available. To do so, an appropriate mathematical transformation of barcodes should be defined, based on the experimentally observed differences between control and unhealthy dendritic shapes. The identified mathematical transformation should be applied to the persistence barcodes that set the synthesis input. This process results in synthesized neuronal morphologies that approximate a target reconstructed population of neurons.

Abnormal dendritic morphology has been linked to brain disorders such as mental retardation (Kaufmann and Moser 2000), schizophrenia (Glausier and Lewis. 2013), autism (Phillips and Pozzo-Miller 2015), and stress disorders (Shansky and Morrison 2009, Dioli et al. 2019, Sandini et al. 2020). We demonstrate how alterations of dendritic shapes that have been associated to mental disorders, such as stress and PTSD (Curran et al. 2017, Dioli et al. 2019, Tornese et al. 2019, Sandini et al. 2020) affect the connectivity of cortical circuits. Even though our digital networks do not yet consist of multiple brain regions, and therefore cannot reproduce the exact medical conditions reported in the literature, we focus of the the effects of local dendritic alterations on a cortical microcircuit. This analysis serves as the first step to link local morphological alterations to whole brain networks and study how “small” changes impact the brain functionality. We simulated two structural changes that are relevant for mental disorders (Curran et al. 2017, Dioli et al. 2019, Tornese et al. 2019): shrinkage of dendritic processes and loss of dendritic branches, by applying two types of mathematical transformations to all the cells within a rodent cortical column. Surprisingly, we discovered that these two types of local dendritic alterations have distinct effects on the resulting cortical networks. Neuronal networks formed from dendrites that are gradually shortened collapse rapidly, as they lose connections almost linearly as a function of the total dendritic extent lost. Networks based on dendrites that lose smaller branches of increasing length are more resilient to connectivity loss. In fact, the effect of these local dendritic changes on the resulting network is observable only when larger branches (≈ 200 *μm*) are lost. This method enables the investigation of the impact of structural neuronal abnormalities and could lead to more advanced diagnostic or treatment techniques.

## Discussion

Since the study of the microscopic structure of the brain beginning in the late 19th century by anatomists such as Deiters, His and Golgi that led to the beautiful and detailed formal descriptions of neuronal morphologies by Ramon y Cajal, there have been numerous astounding breakthroughs in our understanding of the brain. Discoveries range through multiple spatial scales from the dynamical properties of ion channels (Ranjan et al. 2019) and single cell morphologies, physiology and transcriptomics (Janelia, Winnubst et al. 2019, AllenBrain, Gouwens et al. 2019) to the structural and functional connectivity of the whole brain (Wang et al. 2015, Hahn et al. 2019). Due to rapid experimental progress, the burning challenge of our time is assembling all the gathered data into a realistic description of the brain. A fundamental step towards understanding brain function is to elucidate the roles of its fundamental cellular components, primarily the neurons. However, much remains unknown concerning the structural properties of neurons and how the morphology of neurons influences the structural and functional properties of brain networks. A promising approach to discerning the roles of individual neurons in the brain consists of computationally recreating them, i.e., synthesizing them, in order to study their behavior within digital brain networks.

Due to the complex biological growth mechanisms of neurons that result in intricate branching structures, the highly correlated morphological features of neurons are difficult to reproduce. In this study, the computational synthesis of neuronal morphologies was based on the Topological Morphology Descriptor (TMD, Kanari et al. 2018), which retains sufficient information about both the topology and the geometry of a neuronal tree to reproduce the shapes of reconstructed neurons. We proved that the topology-based synthesis preserves correlations between morphological features. Another challenge for synthesis is the sparsity of data for many cell types in the cortex, which makes it difficult to approximate the biological diversity. The minimum number of reconstructed neurons of any particular type sufficient to reproduce the morphological diversity of a population of such neurons is 15 to 20 using our TNS algorithm, which is a relatively small number compared to the thousands of morphologies that are expressed by each neuron type within a brain region. Our algorithm thus overcomes major limitations of previous synthesis techniques, enabling the large scale reconstruction of unique neuronal morphologies to populate large-scale biophysically detailed neuronal networks.

Taking this work a step further, we generalized the TNS algorithm to reproduce cells with altered structural properties, by applying an appropriate transformation to the TMD of reconstructed neurons that are used as input to the synthesis algorithm. In this way, transformation of the TMD enables the study of pathologies, as well. By simulating the effects of stress on single neurons, we demonstrated that the degree and type of degeneration of dendrites influences the nature of global defects exhibited by cortical microcircuits. The next step towards the simulation of brain diseases is to generalize these results to whole brain regions, e.g., the neocortex, thalamus and hippocampus, and to study how different combinations of neuronal deformations can lead to disruptions of brain networks both structurally and functionally. Recent datasets that record the transcriptomics, the electrical and morphologies of cells (Hodge et al. 2019, Gouwens et al. 2019) will be essential for this effort, by enabling the reconstruction of whole brain areas based on their genetic profiles.

While the synthesis of dendrites is already an important step towards the digital reconstruction of more realistic brain networks, the ability to synthesize axonal trees is the next challenge that should be addressed. This task is of particular interest for the computational modeling of brain networks (Wang et al. 2015), as the branching structure of axonal morphologies is an essential determinant of the functionality of a network, by providing the contact points between neurons, and thus defines the connectivity of the network (Van Pelt et al. 2010). In addition, because of their highly complex branching structures, the reconstruction of axons requires considerably more effort and time than dendrites. As a result, only a small number of intact (not cut) axonal reconstructions are available. Spatially embedded synthesis will improve the generation of complex branching patterns, such as cortical axons of both interneurons and pyramidal cells, glial cells, and long-range projecting cells, such as nigrostriatal dopaminergic neurons (Matsuda et al. 2009) and densely connected claustrum cells (Torgerson et al. 2015), and thus allow the digital reconstruction of multiple brain areas and their respective connections.

## STAR Methods

The morphological development of neurons in the brain is a complicated process that depends on both genetic and environmental components (Ledda and Paratcha 2017). The processes that contribute to neuronal growth differ between species, brain regions, and morphological types. Advances in experiments and mathematical and computational models have converged on a set of commonly accepted stages of morphological growth: the initiation of neurites, neurite elongation, axon path-finding, and neurite branching (Graham and van Ooyen 2006). These growth stages are useful for computational modeling of the generation of synthesized neurons. In this study we focus on the computational synthesis of dendrites and thus will not consider axon path-finding. While biological development is not simulated, biological principles of morphological growth inform the design of our computational algorithm that synthesizes dendritic morphologies.

### A. Dendritic synthesis algorithm

The TNS algorithm consists of three main components (Figure 1A): the initiation (section I. Initiation of neurites), branching (section II. Bifurcation / Termination), and elongation (section III. Elongation of neurite) of neurites. The first part of a neuron to be generated is the cell body, i.e., the soma, which is modeled as a sphere (Figure 1A-I), whose radius is sampled from a biological distribution (see SI:Topological Neuron Synthesis algorithms, Algorithm 2). The number of neurites is then sampled from the biological distribution of the corresponding cell type. Each neurite is initialized with a “trunk”, the initial branch of the tree (Figure 1A) and a barcode sampled from set of reconstructed dendrites of the corresponding m-type.

Subsequent steps of the growth take place in a loop. Each branch of the tree is elongated as a directed random walk (Aslangul et al.1993) with memory (see SI:Topological Neuron Synthesis algorithms, Algorithm 3, Figure 1A-III). At each step, a growing tip is assigned probabilities to bifurcate, to terminate, or to continue that depend on the path distance from the soma and are defined by the bars of the selected barcode (Figure 1A-II, see SI:Topological Neuron Synthesis algorithms, Algorithm 3). Once a bar is used, it is removed from the barcode. The growth terminates when all the bars of the input barcode have been used. As an independent step, the diameters of the tree are assigned based on diameter distributions sampled from the biological reconstructions (Figure 1A-IV, section IV. Generation of tree tapering).

#### I. Initiation of neurites

Previous studies have disregarded the direction of the neurite protrusion from the soma despite its importance (Graham and van Ooyen 2006). For some neurites the initial direction is trivially defined; for example cortical apical dendrites typically grow towards the pia. By contrast, the outgrowth direction of basal dendrites superficially appears random and is frequently assumed to be so. An in-depth analysis reveals, however, that this assumption is inaccurate, since the orientations of a neuron’s processes are correlated (see SI, Figure S2). This correlation is captured in the pairwise trunk-angle distribution, which depends on the morphological type, and is used for the initiation of neurites on the soma surface (see SI:Topological Neuron Synthesis Algorithms, Algorithm 3).

For the basal dendrites, the initial point of a single neurite is randomly sampled on the soma surface, then the other dendrites are added successively in places that respect the pairwise trunk-angle distribution. Each neurite trunk consists of a point on the soma surface and an initial direction that is normal to the soma surface. The diameters are independently corrected in the final step of the synthesis algorithm (Figure 1A-IV). The positions of the trunks define the soma shape; the pyramidal soma of excitatory cells originates from the apical dendrite that points towards the pia, while the spherical soma of interneurons arises from the homogeneous positioning of trunks on the surface of their cell bodies.

#### II. Bifurcation / Termination

The neuronal branching process in the TNS algorithm is based on the concept of a Galton-Watson tree (Galton and Watson 1875), which is a discrete random tree generated by the following process. At each step, a number of offspring is independently sampled from a distribution. A neuronal tree consists only of bifurcations, terminations, and continuations, so the accepted values for the number of offspring are: zero (termination), one (continuation), or two (bifurcation). Since the Galton-Watson tree only generates the branching structure and ignores the embedding in space, we modify the traditional process to introduce a dependency of the neuronal growth on the embedding, so that the bifurcation/termination probabilities depend on the path distance of the growing tip from the soma.

Each growing tip is assigned a bar *bar*_*i*_, sampled from the barcode, that includes a starting path distance *b*_*i*_, an ending path distance *d*_*i*_ and a bifurcation angle *a*_*i*_ (see SI:Topological Neuron Synthesis algorithms). At each step the growing tip first checks the probability to bifurcate, then the probability to terminate. If the growing tip does not bifurcate or terminate, then the branch continues to elongate. The probability to bifurcate depends on the starting path distance *b*_*i*_. As the growing tip gets closer to the path distance *b*_*i*_, the probability to bifurcate increases exponentially until it reaches the highest possible value (1.0). Similarly, the probability to terminate depends exponentially on the ending path distance *d*_*i*_.

The probabilities to bifurcate and terminate are sampled from an exponential distribution *e^−λx^*, whose free parameter λ should be wisely chosen. A very steep exponential distribution (high value of λ) will result in cells that are very close to the input and thus will reduce the variance of the synthesized cells. On the other hand, a very low value of λ will result in cells that are almost random, since the dependence on the input persistence barcodes will be decreased significantly. The value of the parameter λ should be of the order of the step size (see SI: Branching-Termination). As a result, we select a critical correlation length λ ≈ 1, so that the bifurcation and termination points are stochastically chosen but are strongly correlated with the input persistence barcodes (See SI: Branching-Termination).

Other synthesis algorithms (Burke et al. 1992, Koene et al. 2009) sample the branching and termination probabilities from independent distributions. In TNS the correlation of these probabilities is captured in the structure of the barcode. When the growing tip bifurcates, the corresponding bar is removed from the input TMD to exclude re-sampling of the same conditional probability. This keeps a record of the neuronal growth history and is essential for reproducing the branching structure. In the event of a termination, the growing tip is deactivated, and the bar that corresponds to this termination point is similarly removed from the input TMD.

At a bifurcation, two new branches are generated (SI:Topological Neuron Synthesis algorithms, Algorithm 4), and the directions of the daughter branches depend on the bifurcation angle *a*_*i*_. Depending on the neurite type, different rules are optimal for the bifurcation angles. For basal dendrites, the optimal rule for bifurcation is to follow the biological bifurcation angle distribution. The apical tree is separated into two parts: the apical tuft, which is the densely branched subtree that is proximal to the cortical surface, and the obliques, which are the shorter branches that emerge closer to the soma. The apical tuft is separated from the obliques by the “apical point”. This point can be accurately identified based on the persistence barcode of the apical tree, as the distance that maximizes the separation between the two modes of the bars distribution. For the apical dendrites, different branching behaviors need to be adopted for the tuft and the obliques. Before the apical point, one of the branches, the major branch, follows the targeting direction (usually the orientation towards the pia). Once the apical point is reached, the apical tufts bifurcate according to the distribution of the biological bifurcation angles.

#### III. Elongation of neurite

A segment is defined as a pair of consecutive points in the neuronal tree that determine a vector of length *L* and with direction *D*_*segment*_, specified by a unit vector. Each synthesized neurite is grown segment by segment. The direction of the segment is a weighted sum of three unit vector terms: the cumulative memory of the directions of previous segments within a branch *M*, a target vector *T*, and a random vector *R* (Koene et al. 2009). The memory term is a weighted sum of the previous directions of the branch, with the weights decreasing with distance from the tip. Different weight functions were tested, but as long as the memory function decreases-faster than linearly - with the distance from the growing tip, its exact form is not relevant. The target vector is defined at the beginning of each branch and depends on the biological branch angles (see SI:Topological Neuron Synthesis algorithms, Algorithm 3). The random component is a vector of fixed length sampled uniformly from three-dimensional space at each step. For computational efficiency the growth of each branch is independent of that of other branches. The tortuosity of the path is defined by three parameters:

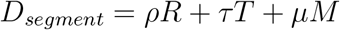
where *ρ* + *τ* + *μ* = 1.

An increase of the randomness weight *ρ* results in a highly tortuous branch, approaching the limit of a simple random walk when *ρ* = 1 (Pearson et al. 1905). If the targeting weight *τ* = 1, the branch will be a straight line in the target direction. Different combinations of the three parameters (*τ, ρ, μ*) can generate more or less meandering branches and can reproduce the large diversity of dendritic sections (see SI:Section elongation).

#### IV. Generation of tree tapering

The thickness of a neuron’s branches should also be accurately reproduced, since thickness is as important as the branching structure for the functional role of neurons (Cuntz et al. 2007, Koene et al. 2009, van Elburg and van Ooyen 2010, Bird and Cuntz 2016). Despite recent great progress in imaging techniques that enables the generation of large numbers of reconstructions (Peng 2008, Haberl et al. 2015, Economo et al. 2016), their resolution is still too limited to allow for accurate determination of diameters, which are on the order of a few microns. As a result, accurate diameters must be computationally inferred from sparse datasets of reconstructed cells. Conde-Sousa proposed a method to correct the swelling of the reconstructed diameters (Conde-Sousa et al. 2017) that usually results in lower mean diameters.

In the absence of a curated dataset, the original diameters of the reconstructed cells are used as input for the synthesis algorithm. Basic morphometrics related to the thickness of neurons are extracted to be used as input for the synthesis of denditic thickness (taper rate, termination thickness, and trunk thickness). These values are used to assign diameters independently to each synthesized dendrite.

The algorithm (see SI:Topological Neuron Synthesis algorithms, Algorithm 5) starts from the tips of the tree and assigns diameters to the termination points sampled from the biological distribution. The tree is then traversed from the tips towards the root (post-order), and the diameters are increased according to the sampled taper rate, as long as the new diameter is less than a sampled maximum diameter, which corresponds to the trunk diameter. When the diameters of all the children of a section have been computed, the parent section is assigned a diameter according to the Rall ratio, which is chosen to be 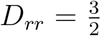 according to literature:

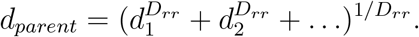

This algorithm results in a distribution of diameters that is statistically similar to the original distribution. In addition, the average diameter of the synthesized cells corresponds to the average biological diameters. The synthesized diameters monotonically decrease with distance from the soma, a property that ensures that biophysical principles of dendrites (Cuntz et al. 2007) are reproduced. Note that the swelling of the dendritic trees, resulting from staining artefacts, is not compensated for and therefore the diameters of the synthesized cells could be overestimated.

### B. Validation process

#### Single-cell validation

In order to validate the quality of single cells and identify individuals of poor quality within the synthesized population, the distributions of key features of each cell are compared against a set of reconstructed cells. To measure a cell’s difference from the reconstructed cells, we compute a statistical normalized distance, which is the difference between the mean value of the test cell *T* and the mean value of the reconstructed population *P*, divided by the standard deviation of the reconstructed population *P*:

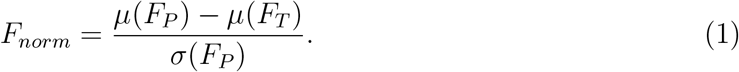

#### Population-to-population morphological validation

Synthesis is validated by comparing the distributions of synthesized cells of a large number of morphological features (see SI:Definition of morpho-electrical terms) to those of the reconstructed cells. Essential features, such as the degree of the dendritic tree (number of terminations), the branch orders and the number of sections, the total length per neurite, and the path length are included in the validation and shown in Figure 4. Each of the morphological distributions of synthesized cells is compared to the reconstructed cells’, using the Kolmogorov-Smirnov distance, which quantifies the dissimilarity between two distributions. The K-S metric measures the maximum distance between two cumulative distributions, ranging from 0 for identical distributions to 1 in the case of maximal difference between them.

#### Electrical simulation of synthesized cells

A biophysically detailed electrical model (e-model) for a L3 TPC was applied in the synthesized morphologies to assess how well the electrical behavior generated by the synthesized morphologies compares with their reconstructed counterparts. The original e-model was obtained by applying a multi-objective optimization of the electrical parameters as described in Markram et al. 2005 to a reconstructed morphology. The e-model has 31 parameters that are used to control the maximal conductances of the ion channels in four morphological areas (somatic, axonal, basal, and apical) and the calcium dynamics and the decay constant of Na channels along the dendrite. Note that in the case of synthesized cells, only dendrites are computationally generated; the axonal morphology is copied from a reconstructed cell. The e-model consists of Hodgkin-Huxley-based channel models for persistent and transient Na/K, high- and low-voltage activated Ca, Kv3.1, Ih and SK calcium-activated potassium channels. When the e-model is instantiated, the axon is replaced by a shorter axon initial segment with diameters based on the original morphology. The constraints consist of electrical features extracted (eFEL) from somatic whole-cell current clamp recordings and dendritic back-propagating action potential features obtained from literature. The stimulation currents used in the experiments and models are scaled by the spiking threshold currents of the cells. The e-model was applied to the synthesized morphologies and, as in the morphological validation, the Fnorm was used to quantify how well the resulting morpho-electrical combination matches with the statistics of the original experimental data.

### C. Synthesis input

Neuronal reconstructions of a large number of cortical morphological classes were used as input to the synthesis algorithm. Biological reconstructions from the BBP dataset of all rodent cortical cell types reported in previous publications (Markram et al. 2015, Kanari et al. 2019) were used as synthesis input. The BBP dataset was chosen as input because of the consistency of the quality of the input morphologies, as they were all generated using the same reconstruction protocol. In addition, the electrical models that have been generated for all cell types (Markram et al. 2015, Van Geit et al. 2016) make the comparison of multiple properties (morphological, electrical) feasible.

A few modifications were performed on the original reconstructions to compensate for reconstruction artefacts. For example, the slicing of the brain tissue and the filling of the cells with biocytin (Horikawa and Armstrong 1988) result in their shrinkage. This effect modifies the tortuosity of the reconstructions (as cells appear more tortuous than they originally were), and the extent of their processes decreases. To compensate for these artefacts, the cells that are used as input for synthesis are initially “;unraveled”, as described in (Markram et al. 2015). Another important artefact is the loss of arborization, due to slicing of the tissue during the reconstruction process. This error is compensated for with a “repair” process described in Markram et al. 2015. Because repair modifies the branching properties of the tree, only cells that have been unraveled, but not repaired, are used as input to the TNS algorithm for the current study. To compensate for the loss of arborizations, trees that contain fewer than three sections are considered cut and thus discarded from the synthesis input during preprocessing of the input data.

## Author contributions

L.K. conceived and implemented the topological synthesis algorithm and performed the computational experiments on the synthesized cells. A.C. implemented and performed morphological validations. W.V.G. developed the software and performed the electrical validation of the synthesized cells. B.C. provided engineering support for the synthesis and validation software. H.D. generated and analyzed the neuronal networks. J.S. contributed to the development of the synthesis algorithms. K.H. and H.M. supervised and discussed the algorithms and results at all stages of this work. All authors discussed the results and commented on the manuscript at all stages.

## Acknowledgments

We kindly thank LNMC lab (EPFL) for the biological reconstructions that were used in this study. We acknowledge Joseph Graham and Guy Atenekeng for crucial input on synthesis algorithm, at the early stages of this work. We thank Jay Coggan for helpful conversations in various stages of this research and for a thorough read of the manuscript. We thank Michael Reimann for his input on the connectivity analysis, Rembrandt Bakker (Bakker et al. 2017) for visualization tools, Nicolas Antille, Cyrille Favreau for visualizations and Caitlin Monney for figure editing and Michael Gevaert for software engineering support.

