## Supplementary material for "Computational synthesis of cortical dendritic morphologies"

### Supplementary Information: Computational synthesis of cortical dendritic morphologies

April 14, 2020

### Topological morphology descriptor algorithm

We briefly explain the TMD algorithm that encodes the branching pattern of a neuronal *tree*, into a persistence barcode TMD(*tree*). , We define a *tree* as an embedding in $3D$-space of a connected, acyclic, directed graph, with a defined root, i.e. the neuronal soma. The details of the algorithm are given in (Kanari et al. 2018). The TMD algorithm encodes the neuronal tree morphology into a topological object that represents its branching pattern. The local fluctuations with minimal content, such as the position of the nodes between branch points, are discarded, and thus the computational complexity of the tree structure is significantly reduced. The TMD algorithm couples the topology of the branching structure with geometric properties of the tree (in this case the radial distance from the soma), encoding its over-all shape into a single descriptor. The algorithm takes as input a set of branch points, or bifurcations, i.e., nodes with more than one child, and leaves, or termination point, i.e., nodes with no children, of the tree and produces a multi-set of intervals, called bars, on the real line, known as a persistence barcode (Ref Carlsson 2009), Fig 1B. Each bar encodes the lifetime of a component in the underlying structure, identifying the radial distance at which a branch is first detected, emerging from a larger subtree (birth, $b_i$) and the radial distance at which the branch terminates (death, $d_i$). Equivalently, the persistence diagram (Fig 1C, Carlsson 2009) represents the pair of birth-death times of each component as a point in the $2D$ plane. Note that all distances are computed in $\mu m$ and for clarity the projection of radial distances is presented in figure Fig 1.

The main concept of the TMD algorithm is presented in Fig 1. It takes as input a rooted *tree* with a real-valued filtration function $f$ defined on the set of the tree's nodes. In this example, the root is the neuronal soma, and the function $f$ is the radial distance from the soma. Each branch within the *tree* (Fig 1A:1-4) is transformed into a bar in the persistence barcode TMD(*tree*) (Fig 1B:1-4) encoding its start (or bifurcation) and end (termination) radial distances. Starting from the terminations, the radial distances of two siblings, i.e., branches starting at the same bifurcation point, are compared. We take as an example branches #2 and #3, which are siblings. The branch with the largest radial distance from the soma, here branch #2, persists, while the smallest sibling #3 "dies" generating a bar within the persistence barcode which encodes the initial and final radial distances $(b_i, d_i)$. Next the remaining branch #2 is compared to the next sibling #4 it encounters following the path towards the root, which is smaller. As a result, a new bar is added to the barcode that corresponds to branch #4 with the respective start and end radial distances. Finally, branch #2 encounters branch #1, which is larger and eventually generates the second longer bar in the barcode. Branch #1, which has the largest termination radial distance, persists all the way to the soma and generates the longest bar, which starts at the soma (radial distance is zero) and ends at the largest radial distance $100\mu m$.

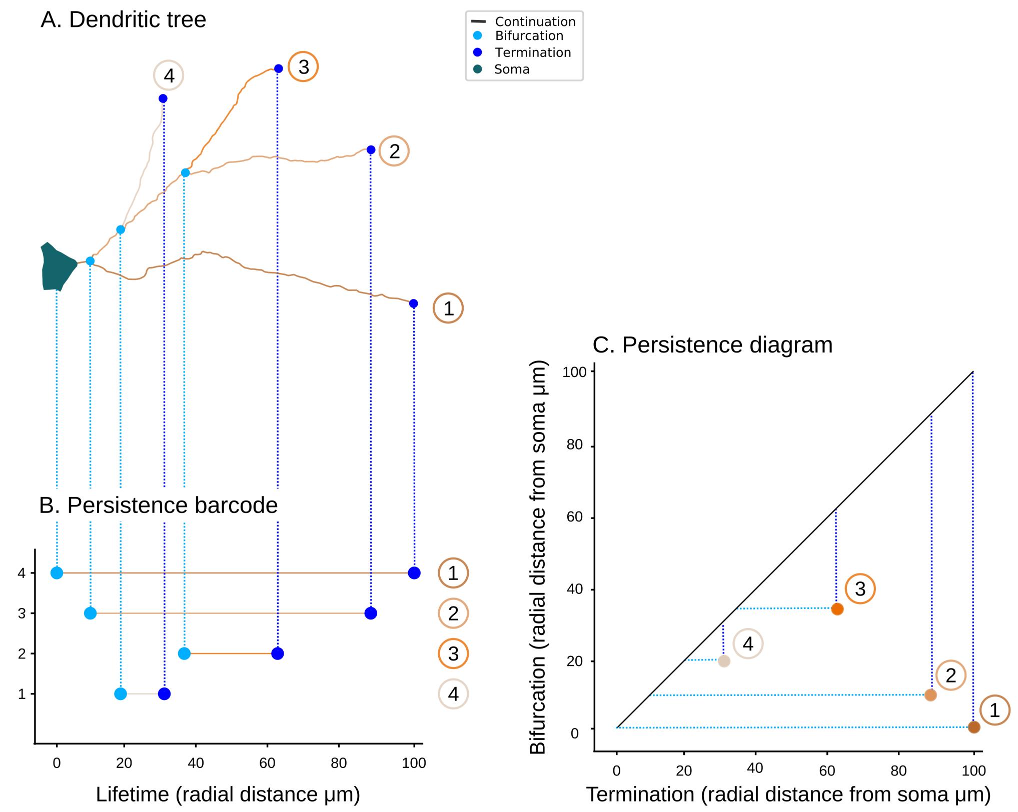


S 1: Illustration of the TMD algorithm. A rooted tree (A) is transformed with the TMD algorithm into the corresponding persistence barcode (B) and persistence diagram (C). The root or soma, colored in green, defines the reference of zero distance (for example, for radial distances). Continuations are presented in black, bifurcations in light blue and terminations in dark blue. The correspondence between the tree (A) and its extracted barcode (B) is indicated by the assignment of the same node numbers for each termination in tree A and each bar in the barcode B. Each bar represents the lifetime of a branch within the tree, defined by a bifurcation and termination radial distance from the soma. The longest bar (1) corresponds to the branch within the tree with the largest termination radial distance. Equivalent to the persistence barcode (B), the persistence diagram (C) represents the start and end radial distances in a $2D$ plane. All radial distances are measured in $\mu m$.

#### Definition of morpho-electrical terms

As a guide to the reader, the main definitions of morphological terms that will be used through this paper are summarized in the following table. Note that these terms might have different meanings elsewhere in the literature. Since morphological terminology is often not consistent through the literature, we clarify the meaning of each term used in this paper.

| Definition of morphological terms |
| --- |
| Soma: the cell body is described as a sphere $S^{c}_{d_s}$ of diameter $d_s$ and center $c_s$. The contour of a soma consists of a set of points on the x-y plane around the soma surface. |
| Neurite: A neuronal tree; a dendrite or an axon. |
| Neurite point: $(x, y, z, d)$, where $(x, y, z)$ are the $3D$ coordinates and $d$ is the dendritic diameter that represents the thickness of the neurite at that point. |
| Branch point: a point in the tree with two (or more) children, also known as a bifurcation point (or mutltifucration if it has more than two children). |
| Neurite tip: a point in the tree without any children, also known as a termination point. |
| Neurite section: a set of points of a neurite, either between two branch points or between a branch point and a termination point. |
| Neurite trunk: the first section of a neurite, starting from the soma and ending at the first branch point. |
| $Vect$: A unit vector in $3D$ space, which is equivalently represented by a pair of angles, and defines a direction, or orientation, in $3D$ space. |

For the definition of morphological features, we use as guide the Petilla terminology paper (features 1-5, Ascoli et al. 2008), as well as some additional features (6-9).

##### Morphometrics glossary

1. Thickness: the dendritic thickness in micrometers.

2. Taper: percentage of thickness narrowing per unit distance.

3. Bifurcation angle: angle between two daughter branches at a branch point (bifurcation).

4. Tortuosity: the straight-line distance between two consecutive branch points divided by the length of the neuronal path between those points.

5. Partition asymmetry: Ratio of the absolute value of the difference and the sum of the number of bifurcations in the two daughter subtrees at a branch point.

6. Radial distance: the end-to-end straight-line distance between the branch point of a section and the soma surface.

7. Path distance: the distance along the neuronal path, between the branch point of a section and the soma surface.

8. Branch order: the number of bifurcations between current position and the root.

9. Section length: total length of a section.

For more details about the electrical features, we refer to (Van Geit et al. 2016). Basic definitions are summarized in the table below.

**Electrical features glossary**

1. AP: action potential.
2. RMP: resting membrane potential.
3. IV: a step protocol used for studying the current-voltage relationship. The currents are kept low to avoid action potential generation.
4. APWaveform: Short step protocol used for studying the action potential shape.
5. Step (120%): a longer (2 seconds) step stimulus, the current amplitude is 120% of the current amplitude that is necessary to trigger one action potential.
6. AHP: after-hyperpolorization after an action potential.
7. ISI: inter-spike interval.

#### Synthesis input

A set of neuronal reconstructions is used as input for the topological neuron synthesis (TNS). The input consists of morphology files (in one of the following formats: neurolucida ASC, SWC, or H5), which contain the three dimensional positions $(x, y, z)$ and the thickness $(d)$ of the neuronal nodes and their adjacency relations, to form a tree.

From the input neuronal morphologies, the persistence barcode of each neuronal tree is generated according to the TMD algorithm described in the previous section. Along with the topological barcodes of the neurons, a set of basic morphometrics, related to the features of the soma and the thickness of the tree, are extracted. Those morphometrics are summarized in the section "Algorithm input distributions". In addition to the biological inputs a set of user-defined parameters, described in "Algorithm input parameters", are also used as input for the TNS algorithm.

#### Biological persistence barcodes

The algorithm that extracts a persistence barcode (Kanari et al. 2018) from a neuronal tree is described in the previous section. The barcodes that are used as input for synthesis are enhanced with the bifurcation angles of their corresponding components, i.e., at each bifurcation point, where a new branch emerges, the bifurcation angle $a$ from its parent is recorded. The $i^{\text{th}}$ bar in TMD($tree$), denoted $bar_i$, corresponds to the $i^{\text{th}}$ branch of the tree and is encoded as a triplet $(b_i, d_i, a_i)$, where $b_i$ is the initial radial distance of the branch, $d_i$ is the terminal radial distance, and $a_i$ is the bifurcation angle at which the component emerges from its parent. Note that $b_i$ and $d_i$ are real numbers, while $a_i$ is a set of four real numbers, which represent the following angles.

- $a_i^1$: angle between child and parent in $x$-$y$ plane
- $a_i^2$: angle between child and parent in $x$-$z$ plane
- $a_i^3$: angle between two children in $x$-$y$ plane
- $a_i^4$: angle between two children in $x$-$z$ plane

In order to recreate a population of biological reconstructions, we need to generate the persistence barcodes for each tree of the reconstructed morphologies. The process of encoding a neuronal tree into a barcode is called "extraction" of the persistence barcodes, and is described in the first section. The complete set of barcodes that will be used as input to the synthesis algorithm is denoted as

$$Barcodes = \{TMD_j | 1 \leq j \leq n_t\},$$

where $n_t$ = # of neuronal trees in the biological population, where for the $j^{\text{th}}$ tree

$$TMD_j = \{bar_i = (b_i, d_i, a_i) | 1 \leq i \leq n_j\},$$

where $n_j$ = # of components (bars) of $TMD_j$.

Barcodes are randomly sampled during the computational growth of a neuron. Each synthesized neurite (dendrite) is generated by a single barcode $TMD$, which corresponds to a unique biological tree.

#### Biological distributions of morphological features

In addition to the persistence barcodes, which encode the topology of the neurites, a number of independent morphometrics need to be measured. The first of these corresponds to the size of the cell body, or soma. The soma is considered to be a sphere and therefore a center and a diameter are sufficient to describe it. The center, or origin of the cell, is a user-defined parameter, while the diameter is sampled from the corresponding biological distribution $D_{ss}$. The number of dendrites $D_{nn}$ and their relative pairwise angles $D_{pa}$ are also sampled from the biological distributions, and specify the number of dendrites that will be generated as well as their intial outward directions. Finally, a set of distributions that describe the thickness of the dendrites is also extracted, sampled from the biological reconstructions. All the morphological features that are used as input to synthesis are summarized in the following table.

| Algorithm input distributions. |
| --- |
| **Soma parameters** |
| $D_{ss}$: Soma diameter. |
| $D_{nn}$: Number of neurites of a specific type within a neuron; basal or apical dendrite. |
| $D_{pa}$: Pairwise angles between neurites, as they emerge from the soma. |
| **Diameter parameters** |
| $D_{tips}$: Diameters of the tips, or terminations of the neurite. |
| $D_{tr}$: Taper rates, define the tapering, i.e., the difference between the diameters at the beginning and end of a section, normalized by the length: $(D_{final} - D_{initial})/length$. The distribution of taper rates is computed for each neuronal section. |
| $D_{rr}$: Rall ratio, which describes the relation between the diameter of a parent and those of its children at a branch point. The value of the Rall ratio is the exponent $n$ such that $D^n = d_1^n + d_2^n + \dots$ at a branch point, where $d_i$ is the diameter of the $i^{\text{th}}$ child and $D$ is the parent diameter. The distribution of Rall ratios is computed for each bifurcation. |
| $D_{md}$: Maximum diameter of each neurite, per type. Under the assumption that dendritic diameter decreases with distance from the soma, this value corresponds to the diameter of the trunk. |

#### Input parameters

The input distributions are computed based on the biological reconstructions. However, the growth process is also controlled by a set of parameters that are defined by the user. The first two parameters, $\tau$ and $\rho$, characterize the elongation of a branch. The origin of the morphology, i.e., the center $c$ of the soma, is an input parameter that allows the user to control the initial position of the cell.

The apical point distance $D_{apical}$ is computed from the persistence barcode, as the point at which a tuft is first encountered. This is a parameter that is not controlled by the user. This distance is used to modify the angles of the bifurcations during the growth of an apical dendrite.

| Algorithm inputs parameters. |
| --- |
| $\tau$: targeting weight that defines how "straight" the generated section will be . |
| $\rho$: randomness weight that defines how "tortuous" the generated section will be. |
| $\mu$: memory weight that defines the independence from previous steps. It is defined by $\mu = 1 - \tau - \rho$. |
| $c$: defines the center of the soma, and the starting point for the growth of the neuron. |
| *step-size*: defines the length of each step during the growth. We represent the step size as a normal distribution with mean $1\mu m$ and standard deviation $0.2\mu m$. |

### Description of synthesis algorithm

#### Soma generation: initiation of neurites

Previous studies have disregarded the direction of the neurite protrusion from the soma despite its importance. In this section, we showcase the correlation of dendritic orientations in reconstructed cells, a property that is essential for the generation of dendrites with biolgically accurate orientations.

The direction of neurite protrusion is particularly important for the generation of dendrites that present an orientation preference (Graham and van Ooyen 2006). For some neurites the initial direction is trivially defined; for example cortical apical dendrites typically grow towards the pia. By contrast, the outgrowth direction of basal dendrites superficially appears random.

In Figure S2 we compare the orientations of reconstructed (A, top, blue), and synthesized (B, middle, red) dendrites to a set of random orientations (C, bottom, yellow). Even though the directions of a population of reconstructed cells visually appear to be random (Figure S2A, left), the pairwise angles between dendrites (Figure S2A, right) present a distinct pattern. Taking into

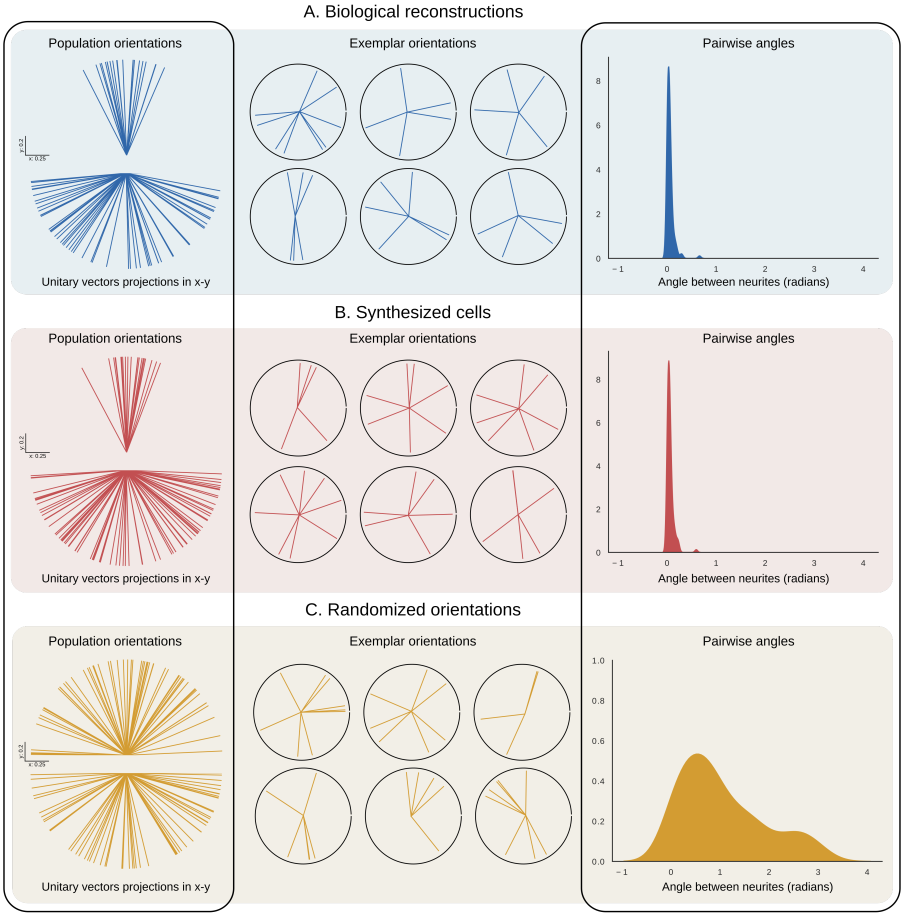


S 2: Orientations of dendrites emerging from somata. The orientation of the reconstructed (A, top, blue) and synthesized (B, middle, red) dendrites are compared to randomly oriented unit vectors (C, bottom, yellow). On the left, orientations of each dendrite within populations of reconstructed (A), synthesized cells (B) and random (C). At the center examples of individual cells from same populations. On the right, the distribution of pairwise angles between all pairs of dendrites of a neuron.

account the pairwise angles of reconstructed dendrites, we can synthesize cells with similar patterns (Figure S2B). However, if the correlations between the dendritic orientations of a neuron are disregarded (Figure S2C), the randomly selected dendritic directions may be either too close to each other or too far away in comparison to the reconstructed cells.

#### Section elongation

A synthesized neurite is grown segment by segment; each segment is assigned a length $L$ and a direction (see Algorithm 3). The direction of a new segment is defined by a unit vector as a weighted sum of three components: the cumulative memory $M$ (see equation 2) of the directions of the previous segments within a branch, a target vector $T$, which is chosen at the beginning of a branch, and a randomly sampled unit vector $R$ (Koene et al. 2009). Therefore, the direction of the new segment is:

$$D_{segment} = \rho * R + \tau * T + \mu * M. \quad (1)$$

The cumulative memory is defined in terms of the last five previous steps of the growth process. If the current segment of the growing branch is the $k^{\text{th}}$ segment, then the five previous steps contribute to the memory as follows:

$$M = \sum_{i=1}^{5} \exp(1 - i) * v_{(k-i)}. \quad (2)$$

The segment length is drawn from a normal distribution with parameters $L_{mean} = 1\mu m$, $L_{std} = 0.2\mu m$. The weights are normalized so that $\tau + \rho + \mu = 1$. Therefore, only two of the input parameters need to be chosen by the user. The different shapes that can be generated by combinations of the weights are presented in Figure S3.

As the randomness weight $\rho$ increases, the branch becomes more tortuous, approaching the limit of a simple random walk (Figure S3, blue, right bottom corner) when $\rho = 1$. On the contrary, an increase of the targeting weight $\tau$ results in straighter branches, which follow the target direction (purple straight line in Figure S3, left down corner). The memory $\mu$ has a more complicated effect on the generated branch. For high values of the targeting parameter $\tau$, the line is already straight, so the effect of memory weights is not significant. For lower values of the targeting parameter $\tau$, the memory decreases the local randomness by preserving directional correlations between consecutive growth steps. As a result, for larger memory weights $\mu$, the branches are more curved at longer distances, but more straight locally.

For a dendrite, the effects of randomness and targeting are more complex. Synthesized cells for a set of varying combinations of randomness and targeting are presented in Figure S4. For smaller targeting ($\tau \approx 0.1 - 0.2$) and larger randomness ($\approx 0.4$) the synthesized dendrites are too short to reach the original dendritic extent, as demonstrated in the corresponding persistence diagrams of radial distances (Figure S4). For a broad range of parameters ($\rho \leq 0.3$ and $0.1 \leq \tau$), the radial distances of synthesized trees

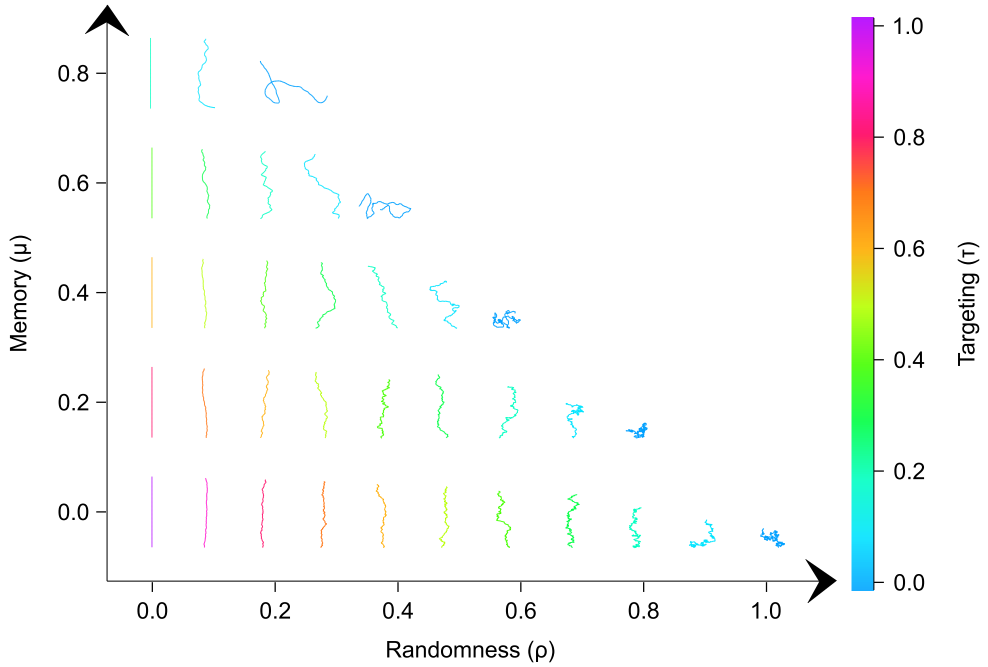


S 3: Synthesized branches for different input parameters. The $y$-axis represents the memory weight $\mu$, the $x$-axis represents the randomness weight $\rho$, and the colorbar corresponds to the targeting weight $\tau$.

(Figure S4, red) approximate accurately the reconstructed cells (Figure S4, blue).

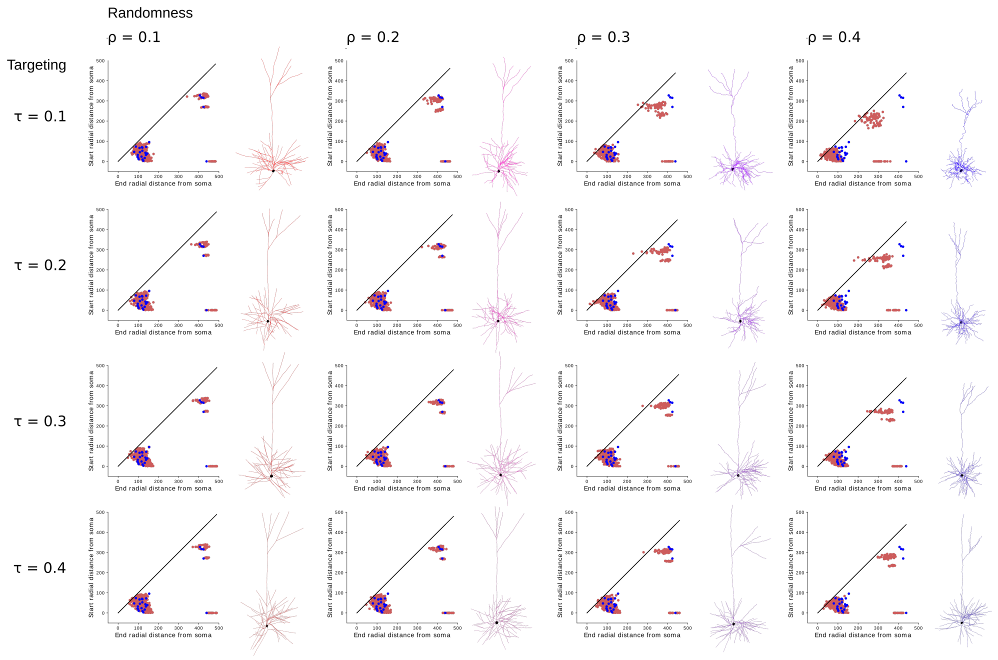


S 4: Sensitivity analysis for elongation parameters. Synthesized dendrites with different input parameters, and the corresponding radial distance persistence diagrams for each tree. Blue diagrams correspond to the target reconstructed cell, red diagrams correspond to synthesized trees for different parameters ($x$-axis corresponds to randomness weight $\rho$, $y$-axis corresponds to targeting weight $\tau$).

#### Branching - Termination

Each growing tree is assigned a barcode TMD. Each growing branch of the tree is assigned a bar $bar_i$, which consists of a start path distance $b_i$, an end path distance $d_i$, and a bifurcation angle $a_i$. The branch will continue to grow until the branch either bifurcates or terminates. To determine whether one of these actions occurs, we compare the path distance of the current position $pd_{tip}$ to $b_i$ and $d_i$ according to the following exponential probability distributions.

$$P_B(\textbf{bifurcation} \mid pd_{tip}) = \exp\left(\lambda * (pd_{tip} - b_i)\right) \tag{3}$$

$$P_T(\textbf{termination} \mid pd_{tip}) = \exp\left(\lambda * (pd_{tip} - d_i)\right) \tag{4}$$

To check if the branch bifurcates, a number, *random*, between 0 and 1 is randomly sampled and compared to the probability of equation (3). If the random number if smaller than this probability, i.e.,

$$random \leq \exp\left(\lambda * (pd_{tip} - b_i)\right), \tag{5}$$

then a bifurcation occurs. If not, another number between 0 and 1 is randomly sampled and compared to the probability of equation (4); if the new random number is less than this probability, then the branch terminates.

If $pd_{tip} < b_i$, the probability to bifurcate $P_B$ is less than 1 and increases as the target value $b_i$ is approached. Once $pd_{tip} = b_i$, $P_B = 1$ and therefore the branch will necessarily bifurcate. The same process holds for the termination of a branch. For very large $b_i$, a bifurcation is very unlikely to occur for small $pd_{tip}$, as $P_B$ is close to 0. In general, if $pd_{tip}$ is much smaller than $b_i$, the probability to bifurcate is very low. Quantitatively, this property depends on the selection of the parameter $\lambda$ of the exponential probability function. The parameter $\lambda$ controls the slope of the probability distribution for bifurcation and termination. The parameter $\lambda$ should be wisely chosen in order to generate biologically relevant neurites (see Figure S5).

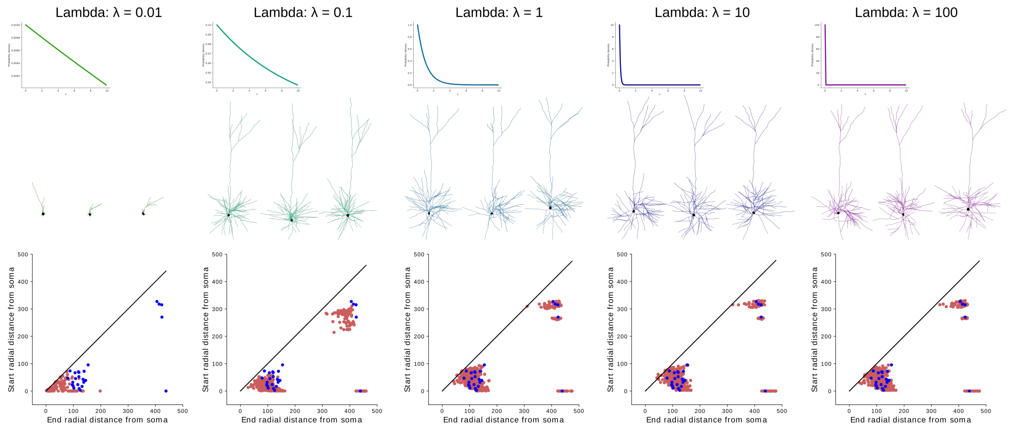


S 5: Sensitivity analysis for the bifurcation/termination parameter $\lambda$. Synthesized dendrites with different input parameters $\lambda$, and the corresponding radial distance persistence diagrams for each tree. Blue diagrams correspond to the target reconstructed cell, red diagrams correspond to synthesized dendrites for $\lambda : 0.01 - 100$ from left to right.

A very steep exponential distribution (high value of $\lambda$) generates cells that are very close to the biological input and thus the variability of the synthesized cells is reduced (right panel Figure S 5). On the other hand, a very low value for $\lambda$ generates cells that are almost random, since the probabilities to bifurcate and terminate are high long before the target values are approached (left panel Figure S 5). If the value $\lambda$ is chosen to be of the order of the step size, the bifurcation will occur within a few steps of the target path distance. As a result, the generated shapes will not be identical to the input tree, thereby increasing the variability of synthesized cells, but will preserve the overall shape of the input tree, generating biologically acceptable branching structures.

In the event of a bifurcation, two new branches are generated (see Algorithm 4), with associated directions depending on the input bifurcation angle $a_i$. Three branching methods were examined: symmetric, biased, and composite. The "symmetric method" imposes that the two daughter branches emerge at the same angle from their parent branch. The bifurcation angle is split in two and equally distributed between the daughter branches. The "biased method" imposes that one of the daughter branches continues to grow towards the direction of its parent and therefore the split is asymmetric. The "composite method" combines the symmetric and the biased methods. For the "composite method" multiple angles are required: the bifurcation angles $a_i^1, a_i^2$ encode the angle between the parent branch and the first child, while the bifurcation angles $a_i^2, a_i^3$ define the angle between the two children branches.

#### Definition of tree thickness

The thickness of a neuron's branches should also be accurately reproduced, since thickness is as important as the branching structure for the functional role of neurons. In the absence of a curated dataset, the original diameters of the reconstructed cells are used as input for the synthesis algorithm. The reconstructions are analyzed with https://github.com/BlueBrain/NeuroMNeuroM to extract the morphometrics related to the thickness of neurons that are used as input for the synthesis of diameters (taper rate within a section, $D_{tr}$, the diameters of the termination, $D_{tips}$, and the trunk diameters, $D_{md}$) of a tree. These values are used to assign diameters independently to each synthesized dendrite.

The algorithm starts from the tips of the tree and assigns diameters to the termination points sampled from the biological distribution $D_{tips}$. The tree is then traversed from the tips towards the root (post-order), and the diameters are increased according to the biologically sampled taper rate $D_{tr}$, as long as the new diameter is less than a sampled maximum diameter, which corresponds to the trunk diameter $D_{md}$. When the diameters of all the children of a section have been computed, the parent section is assigned a diameter according to the Rall ratio, which is chosen to be $D_{rr} = \frac{3}{2}$ according to literature:

$$d_{parent} = (d_1^{D_{rr}} + d_2^{D_{rr}} + \dots)^{1/D_{rr}}.$$

This algorithm results in a distribution of diameters that is statistically similar to the original distribution. In addition, the average diameter of the synthesized cells corresponds to the average biological diameters. The synthesized diameters monotonically decrease with distance from the soma, a property that ensures that basic physical principles of dendrites (Cuntz et al. 2007) are reproduced. Note that the swelling of the dendritic trees, resulting from staining artefacts, is not compensated for and therefore the diameters of the synthesized cells could be overestimated.

### Validation of synthesized basal and apical dendrites

#### Population-to-population validations

We validate the synthesized neurons by comparing the distributions of their morphometric features to those of the biological cell reconstructions. In the following figures, we show the comparisons for the Sholl analysis and for further key features like the degree of the dendritic tree (number of terminations), branch orders, number of sections per neurite, total length, section intermediate and section terminating lengths, and path length to the terminal tips. We report summary statistics like means, standard deviations, medians, and sample sizes, and use Kolmogorov-Smirnov distance to quantify the dissimilarity between the distributions. Concretely, it measures the maximum distance between the two empirical cumulative distributions, ranging from 0 in the case of identical distributions to 1 in the case of maximal difference, for example when the distributions are completely shifted. In the case of discrete distributions like that of the branch orders, we use the adapted version of the Kolmogorov-Smirnov test described in Arnold and Emerson (see reference). Additional feature comparisons between the synthesized and reconstructed cell populations are given in the following table. These include the number of neurites, number of sections per neurite, number of bifurcations, radial (Euclidean) distance, bifurcation angles, tortuosity, and partition (as an expression of the asymmetry of the cells).

### Topological Neuron Synthesis algorithms

The TNS algorithm consists of three main components: the initiation (Algorithm 2), the elongation (see Algorithm 3) and the branching (Algorithm 4) of neurites.

---

**Algorithm 1** Synthesizer

---

**Input:**

$D_{soma} = \{D_{ss}, D_{nn}, D_{pa}\}$ ▷ (see Biological distributions)
$D_{diam} = \{D_{tips}, D_{tr}, D_{rr}, D_{md}\}$ ▷ (see Biological distributions)
$Param = \{c, \tau, \rho, step - size\}$ ▷ (see Input Parameters)
$Barcodes$ ▷ (see Biological barcodes)
**function** SAMPLE(distr) := draws from input distribution

Generate a Soma and Neurites using (Alg 2, $D_{soma}$, $c$) ▷ (each neurite is initialized with a point on the soma surface, which also defines an initial direction $dir_1$)
**for** neurite in Neurites **do**
    $TMD$ = SAMPLE($Barcodes$)
    Sort $bars$ in $TMD$ from longest to shortest
    Initialize first section $sec_1$ with the longest $bar_1$
    $Active \leftarrow sec_1$
    **while** $Active$ sections **do**
        **for** Section $sec_k$ in $Active$ **do** ▷ (a section gets a target direction $dir_k$ and a bar $bar_k$)
            Grow a section using (Alg 3, $dir_k$, $bar_k = (b_k, d_k, a_k)$)
            Remove $bar_k$ from $TMD$ ▷ (each $bar$ can be used only once)
            **if** $status = Bifurcate$ **then**
                Generate children using (Alg 4, $TMD, bar_k, dir_k$)
                Add children to $Active$ sections
            **else if** $status = Terminate$ **then**
                Section growth terminates

            Remove current section $sec_k$ from $Active$

Generate diameters using (Alg 5, $D_{diam}$)

**Output:**

A neuron: set of points, diameters and their connectivity.

---

The first part of a neuron to be generated is the cell body, i.e., the soma (see Soma generation: initiation of neurites), whose radius is sampled from a biological distribution. The number and the orientation of the neurites are sampled from the biological distributions.

---

**Algorithm 2** Generate soma - neurites initialization

---

**Input:**

$c, D_{soma} = \{D_{ss}, D_{nn}, D_{pa}\}$ ▷ (see Biological distributions)
**function** SAMPLE(distr) := draws from input distribution
$d_s = \text{SAMPLE}(D_{ss})$ ▷ (soma size)
*Soma* is a sphere of diameter $d_s$ and center $c$: $S^c_{d_s}$
$nn = \text{SAMPLE}(D_{nn})$ ▷ (number of neurites)
Sample random unit vector $Vect_1$ of neurite $N_1$ on $S^c_{d_s}$ surface.
First point of $N_1$ is $x^1_1 = c + Vect_1$
**for** Neurite $(N_i|2 \leq i \leq nn)$ **do**
    $Vect_i = Vect_{i-1} + \text{SAMPLE}(D_{pa})$
    First point of $N_i$ is ${x_1}^i = c + Vect_i$

**Output:**

A contour of soma points on $S^c_{d_s}$
Initial points $x^i_1$ for each neurite $N_i$

---

A segment is defined by a length $L$ and a direction $v_k$, specified by a unit vector. The direction of the segment is a weighted sum of three unit vector terms: the cumulative memory of the directions of previous segments within a branch $M$, a target vector $T$, and a random vector $R$.

---

**Algorithm 3** Elongate section

---

**Input:**
  $\tau, \rho, dir$, $bar_k = (b_k, d_k, a_k)$, $x_0$
  $\mu = 1 - \tau - \rho$   ▷ (Normalization of weights to 1)
  **function** PD(point) := path distance of point from soma surface
  $n = 1$
  $status = Continue$
  **while** $status$ is $Continue$ **do**
    $L$: length, $R$: unit vector randomly sampled
    $M = \sum_{i=1}^{5} \exp(1 - i) * v_{(k-i)}$
    $v_k = \rho * M + \tau * T + \mu * M$
    $x_{n+1} = x_n + L * v_k$
    $rand \in [0, 1)$ randomly sampled
    **if** $rand \leq \exp\left(\lambda * (\mathrm{PD}(x_{n+1}) - b_k)\right)$ **then**
      $status = Bifurcate$
    **else if** $rand \leq \exp\left(\lambda * (\mathrm{PD}(x_{n+1}) - d_k)\right)$ **then**
      $status = Terminate$
    **else**
      $status = Continue$
**Output:**
    Section points and status: $Bifurcate$ or $Terminate$.

---

Each growing tip is assigned a bar $bar_i$, sampled from the barcode, that includes a starting path distance $b_i$, an ending path distance $d_i$ and a bifurcation angle $a_i$.

At a bifurcation, two new branches are generated, and the directions of the daughter branches depend on the bifurcation angle $a_i$.

**Algorithm 4** Bifurcate

**Input:**

TMD, $dir_k$, $bar_k = (b_k, d_k, a_k)$

**function** SPLIT(vect, $a$) := returns two unit vectors $dir_1, dir_2$ according to the input vect and a set of angles $a$

▷ (see section Branching - Termination)

$dir_1, dir_2 = \text{SPLIT}(dir_k, a_k)$

Find next available indices $i$ in $TMD$ for which $min(b_i)$ and $d_i \leq d_k$

Generate child $sec_1$: $\leftarrow x_n^k, dir_1, bar_1 = (b_i, d_k, a_i)$

Generate child $sec_2$: $\leftarrow x_n^k, dir_2, bar_2 = (b_i, d_i, a_i)$

**Output:**

Two new sections, each initialized with $x_0$, $dir$ and $bar$.

**Algorithm 5** Diametrizer

**Input:**

$D_{diam} = \{D_{tips}, D_{tr}, D_{rr}, D_{md}\}$ ▷ (see Biological distributions)

**function** SAMPLE(distr) := draws number from distribution

**for all** Neurite tips **do** $d_{tip} = \text{SAMPLE}(D_{tips})$

*Active* ← *tips*

**while** Active **do**

  **for** Section in Active **do**

    $taper = \text{SAMPLE}(D_{tr})$

    **for** Point in Section **do** ▷ From termination to the root

      $d_{new} = d_{n+1} + taper * length$

      **if** $d_{new} \leq D_{md}$ **then**

        $d_n = d_{new}$

      **else**

        $d_n = d_{n+1}$

    Remove Section from Active

    **if** all siblings $1, 2, \ldots$ computed **then**

      $n = \text{SAMPLE}(D_{rr})$

      $D_{parent} = (d_1^n + d_2^n + \ldots)^{(1/n)}$

      Add parent Section to Active

**Output:**

Assigns new values to the diameters of the neurites.

### Results

#### Identifying Outliers in the Population of Synthesized Cells

As an additional validation of synthesized cells, we developed a method to identify outliers in the population of synthesized neurons, by comparing each synthesized cell to the population of reconstructed cells. The percentage of detected outliers in the synthesized population gives a measure of accuracy of the synthesis process.

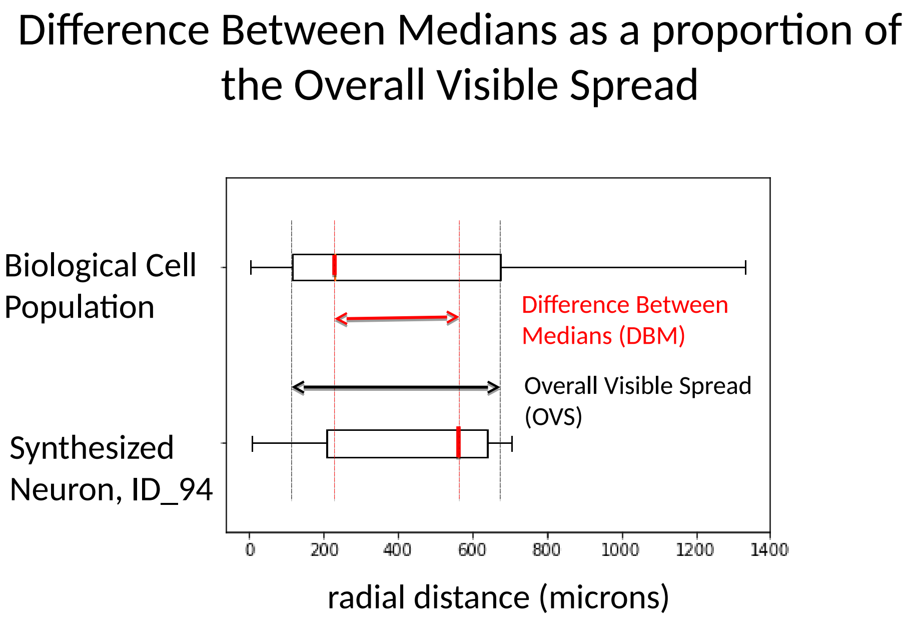


S 6: Statistical measurement for the identification of outliers: the absolute value of the Difference Between the Medians divided by the Overall Visible Spread (DBM/OVS).

To identify outliers, we compare the distributions of key features of each neuron, such as section lengths, bifurcation angles etc, to the reconstructed population. We quantify the distance between the distributions with the ratio of the absolute value of the Difference Between the Medians divided the Overall Visible Spread (DBM/OVS, see Figure S 6). This is an intuitive measure of the difference between the medians of the two distributions with respect to their joint dispersion. The Overall Visible Spread is usually defined as the range between the minimum 25th percentile of the two distributions and the maximum 75th percentile of the distributions. Its minimum is 0 when the medians of the distributions coincide, and the two distributions are very close to each other. Its maximal value is 1 in the special case where the smaller median coincides with the smaller 25th percentile and the larger median with the largest 75th percentile. The closer the DBM/OVS gets to 1, the larger is the difference of medians with respect to the overall spread of the distributions, making it likely to reject the neuron in question. We modify this definition due to the large variance of features within the set of reconstructed cells, and use the OVS defined as the range between the 10th to 90th percentile of the population.

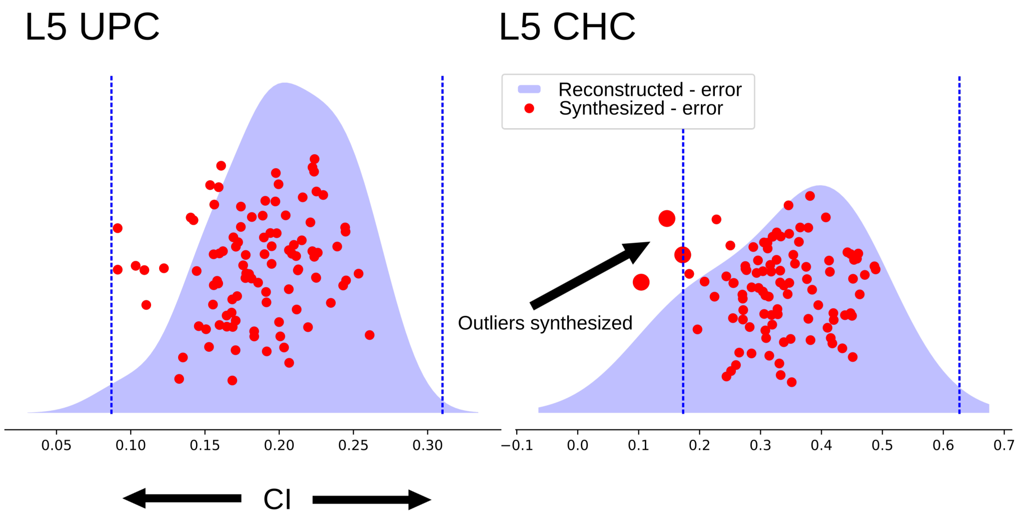


S 7: Detection of outliers in synthesized cells. For two population of cells (layer 5 UPC and CHC), the distribution of DBM/OVS for reconstructed cells is shown in blue, and the corresponding DBM/OVS of synthesized cells in red dots. The dotted lines define the OVS (for 90th precentile) of the distribution. Anything that is outside the OVS is considered an outlier.

The DBM/OVS measure works better than other measures of standardized difference of means, such as Hedges' g, for non-symmetric distributions (see two last references). A cell is considered an outlier when at least one of the key features mentioned above is outside of certain feature-specific thresholds. We choose the thresholds so that the reconstructed biological cells are not rejected as outliers, since they represent the gold standard. One can imagine the thresholds as defining a hypercube in the space of features. If the feature vector of a synthesized cell falls out of this cube, it is rejected as an outlier.

We synthesized 100 cells for each m-type and computed the number of outliers based on the DBM/OVS for the $10 - 90\%$ percentile of the input population. In the following tables (Table 1: PC, Table 2: Interneurons) we report the number of outliers per 100 synthesized cells, and the number of input cells. Note that we report only cortical m-types for which at least three biological reconstructions were available.

As an example, we provide the results for a zero-outlier m-type (L5_UPC) and for one with outliers (L5_CHC) in Figure 7. The population of layer 5 UPC cells has a large number of reconstructed cells (28) and therefore the DBM/OVS of synthesized cells are within the limits of the reconstructed ones. On the contrary, layer 5 CHC consists of only three reconstructed cells, and thus the number of outliers for this m-type is much larger. In addition, the spread of DBM/OVS for the L5_UPC is much smaller than the L5_CHC. A large number of available reconstructions is clearly essential for the accurate definition of the synthesis input.

| | Pyramidal cells | |
|---|---|---|
| M-type | Number of outliers | Number of cells |
| L2_IPC | 6 | 4 |
| L2_TPC:A | 0 | 4 |
| L2_TPC:B | 0 | 33 |
| L3_TPC:A | 0 | 46 |
| L3_TPC:B | 0 | 17 |
| L4_SSC | 0 | 10 |
| L4_TPC | 0 | 37 |
| L4_UPC | 0 | 33 |
| L5_TPC:A | 0 | 72 |
| L5_TPC:B | 0 | 42 |
| L5_TPC:C | 0 | 25 |
| L5_UPC | 0 | 28 |
| L6_BPC | 0 | 25 |
| L6_HPC | 0 | 23 |
| L6_IPC | 1 | 27 |
| L6_TPC:A | 0 | 25 |
| L6_TPC:C | 0 | 19 |
| L6_UPC | 0 | 17 |

| | Interneurons | |
|---|---|---|
| M-type | Number of outliers | Number of cells |
| L1_DAC | 5 | 16 |
| L1_HAC | 4 | 27 |
| L1_LAC | 2 | 10 |
| L1_NGC-DA | 0 | 19 |
| L1_NGC-SA | 7 | 19 |
| L1_SAC | 2 | 14 |
| L23_BTC | 0 | 14 |
| L23_CHC | 0 | 10 |
| L23_DBC | 1 | 18 |
| L23_LBC | 0 | 41 |
| L23_MC | 3 | 29 |
| L23_NBC | 0 | 18 |
| L23_NGC | 1 | 4 |
| L23_SBC | 0 | 22 |
| L4_BTC | 4 | 7 |
| L4_DBC | 4 | 7 |
| L4_LBC | 0 | 31 |
| L4_MC | 0 | 17 |
| L4_NBC | 0 | 25 |
| L4_SBC | 0 | 14 |
| L5_BP | 0 | 3 |
| L5_BTC | 1 | 11 |
| L5_CHC | 16 | 3 |
| L5_DBC | 3 | 9 |
| L5_LBC | 1 | 21 |
| L5_MC | 0 | 38 |
| L5_NBC | 0 | 22 |
| L5_SBC | 4 | 4 |
| L6_LBC | 0 | 24 |
| L6_MC | 0 | 20 |
| L6_NBC | 0 | 11 |
| L6_SBC | 2 | 5 |

#### Marginal probabilities synthesis

We modify the synthesis algorithm to investigate the impact of using marginal branching probabilities instead of joint branching probabilities through persistence barcodes. To do this, we use the information encoded in the persistence barcode ($b_i$, $d_i$, $a_i$) as independent variables. First, we synthesize cells by decoupling the end and start of each bar within a barcode. Instead of using the linked ($b_i$ - $d_i$) we sample the bifurcation from a set of all $b_i$ and the termination from the set of all $d_i$ (see results in Figure S 8A, III). Finally, we decouple the bifurcation angles $a_i$ from the branching probabilities $b_i$, by sampling a random angle at each bifurcation (see results in Figure S 8A, IV).

The results are presented in Figure S 8. For all cases of decoupled algorithms, both the dendritic shapes (see Figure S 8,III, IV) and the persistence diagrams densities are significantly different from the reconstructed and TNS-synthesized cells (Figure S 8, I, II). However, the marginal probabilities for bifurcation and termination are accurately reproduced (Figure S 8A) for all cases.

We have thus proved that the persistence barcodes encode the relevant information about the biological branching structures required for the accurate generation of dendritic shapes. In addition, the links between the bifurcation, the termination, and the respective bifurcation angles are essential for synthesizing biologically accurate cells. The persistence barcodes are therefore a necessary constraint on the synthesis inputs.

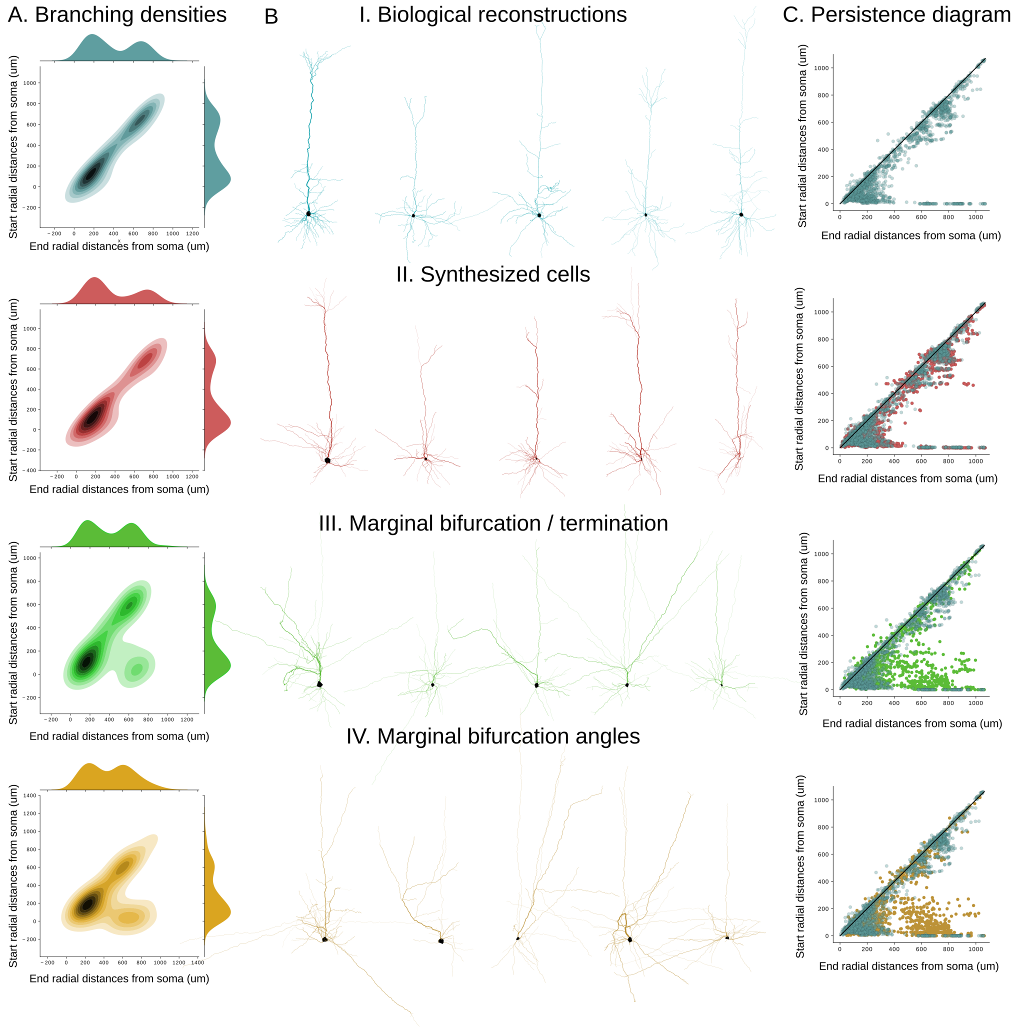


S 8: Comparison of synthesized cells for different synthesis methods. A. Density and marginal projections of persistence diagrams for reconstructed cells (I), synthesized cells (II), synthesized without correlation of branching / termination (III), and synthesized without correlation between branching and bifurcation angles (IV). B. Examples for the same data. C. Respective persistence diagrams.
